# A universal protein ladder for standardisation of diverse FRET assays

**DOI:** 10.64898/2026.03.05.709886

**Authors:** Evelyn R. Smith, Katie L. Gelder, Lewis Hunter-Craig, Daniel A. Bose, Timothy D. Craggs, Alison E. Twelvetrees

## Abstract

Fluorescence resonance energy transfer (FRET) is the highly distance dependent (3-10 nm) transfer of energy from a donor to an acceptor fluorophore, with transfer efficiency inversely proportional to the distance between the fluorophores. Consequently FRET serves as a powerful spectroscopic ruler for probing molecular interactions. Whilst cell based FRET assays report bulk relative changes in FRET efficiency in a population, single molecule FRET (smFRET) is capable of deconvoluting these population averages into distinct structural states. However, the lack of universal benchmarks prevents the direct translation of ***in vitro*** distance measurements to the intracellular environment and vice versa. Here, we present a modular protein ladder designed to harmonize FRET data across diverse platforms. Using an engineered repeating TPR motif and self-labeling enzymes, we demonstrate that our standards yield consistent FRET efficiencies across expression systems (mammalian and bacterial) and labelling strategies (self labelling enzymes and click chemistry with non-canonical amino acids). By providing a predictable calibration curve, the ladder enables interpolation between different experimental FRET modalities, including confocal smFRET, flow cytometry based-FRET and Fluorescence Lifetime Imaging Microscopy FRET (FLIM-FRET). This is the necessary infrastructure to relate molecular distances from the test tube to the cell.

## INTRODUCTION

Fluorescence resonance energy transfer (FRET) occurs between a pair of fluorophores with spectral overlap between the emission spectrum of the donor fluorophore and the excitation spectrum of the acceptor fluorophore. FRET is detected as emission from the acceptor fluorophore upon donor excitation. FRET is highly distance dependent; the efficiency of the energy transfer is inversely proportional to the sixth power of the distance between the two fluorophores. For most fluorophores used in biological systems, FRET only occurs when the two fluorophores are within 10 nm of each other ^1,2^. Fluorophore quenching can occur below 3 nm of separation ^3,4^, giving an effective dynamic range of 3-10 nm when measuring the efficiency, E, of energy transfer. It is possible to determine the distance between the two fluorophores based on E, which offers a resolution beyond the diffraction limit of light and super-resolution microscopy techniques. This has been used to gain detailed structural information through the application of single-molecule FRET (smFRET), particularly for dynamic molecules that are not amenable to more conventional structural biology techniques ^1,5–8^.

As FRET measurements are based on fluorescence, a common tool across disciplines in the biological sciences, they are highly versatile. FRET measurements can be taken on a diverse array of instruments. They include variants of single molecule smFRET using either freely diffusing or surface immobilised systems ^1^ and ensemble measurements, either in solution in fluorescence plate readers or through light microscopy of cells, from widefield to super-resolution modalities ^9^. As the lifetime of fluorophores is impacted by the FRET process, fluorescence lifetime imaging microscopy (FLIM) can also be used to explore FRET in biological systems, with the added advantage of measurements being independent of fluorescence intensity (FRET-FLIM). This breadth of applications also emphasises the untapped potential of FRET measurements to link nanoscale observations across systems.

Despite their potential utility, absolute FRET efficiencies and corresponding accurate distance determination are often technically demanding to calculate. There are several photophysical factors that contribute to this, including: The donor emission spectrum ‘tail’ extending into the spectral window used to detect the acceptor (*α*); The excitation wavelength used for the donor slightly exciting the acceptor directly (*δ*); The overall detection (*γ*) and excitation (*β*) efficiency of donor and acceptor fluorescence rarely being the same in any one acquisition system. Careful calibration creates correction factors for *α*, *β*, *δ*, and *γ* ^10^. However, this is complicated still further in ensemble and cellular systems, where sample heterogeneity (molecules rarely existing in a single, homogenous state) and labelling stoichiometry (the ratio of donor to acceptor molecules may not be 1:1), severely complicates intensity-based quantification. Therefore, many FRET based assays take a ratiometric approach, looking at the relative changes of the ratio of fluorescence intensity between the acceptor and the donor fluorophores sometimes known as FRET indices or relative proximity ratios. These factors combine to create challenges in the reproducibility of both single molecule and ensemble FRET measurements in themselves. They also prevent integrating data across experiments that use FRET as a measurement technique, but differ in sample preparation and class of instrument ^11^.

Due to the technical limitations outlined above, researchers using FRET have historically struggled with apparent FRET efficiency values that vary significantly between instruments and experimental modalities. To address this, in a multi-laboratory comparative blind study ^12^, standardised double-stranded DNA constructs were used in conjunction with benchmarking protocols to demonstrate that instrumental biases can be corrected using alternating-laser excitation (ALEX) to apply fluorophore specific correction factors ^10^. However, while linear DNA provides a rigid and predictable scaffold in vitro, it cannot be used for most in cell experiments due to stability and delivery constraints. Subsequent efforts have used purified proteins as dynamic benchmarks to demonstrate the reproducibility of structural state transitions across labs ^13^. Whilst this has been invaluable for method validation, the measurements cannot be scaled in a linear manner and the maleimide labelling chemistry is not compatible with cellular measurements.

Similarly, polyproline has been used *in vitro* as a spectroscopic ruler ^14^, but such long polyproline peptides (up to 40 residues in length) are still flexible and cause ribosome stalling when translated in cells ^15^. These examples highlight the utility of standard molecules for method development, however their application is currently limited to *in vitro* studies using smFRET. Conversely, control molecules in cellular studies are typically limited to a fusion between the two chosen fluorophores and co-transfected single fluorophores. Whilst providing positive and negative controls respectively, these do not encompass the full dynamic range of FRET. Where linkers are used to capture the FRET dynamic range, these have historically been flexible peptide linkers ^16^. While flexible peptides can successfully decrease FRET efficiency in ensemble measurements ^16,17^, they are less suited to smFRET, as the fluorophores continuously sample the interflurophore distance and thus substantially decrease the signal:noise. Flexible or disordered linkers are also susceptible to structural biases within the cell, for example in response to changes in osmotic pressure, further limiting their broad use for interpolation of data between the test tube and the living cell ^17^. Consequently there is an unmet need for a protein-based reference system that can be labeled site-specifically with donor and acceptor fluorophores and applied effectively in both *in vitro* and in cell experiments.

Here, we address this technological gap by developing a modular protein FRET ladder based on an engineered repeating TPR motif. Our ladder provides a linear scale of predictable FRET efficiencies that remains consistent across expression systems and labeling strategies. This means it can be used both intracellularly and *in vitro* and across FRET imaging modalities. We demonstrate the consistency of our engineered ladder across confocal single molecule FRET (smFRET), flow cytometry based-FRET and fluorescent lifetime FRET (FLIM-FRET). This combination of techniques spans *in vitro* single molecule and intracellular measurements as well as intensity and lifetime-based approaches. Using the same standard ladder across the assays allows the correlation of different FRET based outputs, creating a method for direct translation of *in vitro* smFRET insights into structural dynamics through to in situ measurements in cells.

## RESULTS

### Rational design of a cross-platform protein ladder for FRET

When developing the ladder we had two key design constraints. In order to be usable in a range of applications, fluorescent labelling had to be robust for both *in vitro* purified proteins and in living mammalian cells. We also wanted a modular design that spanned the FRET dynamic range. A protein rather than DNA based ladder is the most suited to diverse expression systems, as small linear DNA fragments are unstable in cells. Consequently, labelling chemistries have to be suitable for living cells whilst still allowing incorporation of both donor and acceptor fluorophores with minimal linkage errors. Similarly, introducing exogenous functional proteins that happen to be of useful dimensions into a living system will have unpredictable impacts on data interpretation.

To alter the distance between the fluorophores in our protein FRET standard ladder, we chose to use an idealised tetratricopeptide repeat (TPR) motif ^18^. The motif, consisting of two antiparallel *α*-helices, occurs naturally in hundreds of different proteins. This idealised form is highly stable and monomeric starting with a three amino acid N-cap and finishing with a solvating helix to improve solubility. The two *α*-helices of the TPR motif can be repeated, creating a modular ladder increasing by two *α*-helices between each step (Figure 1A).

**Figure 1:**
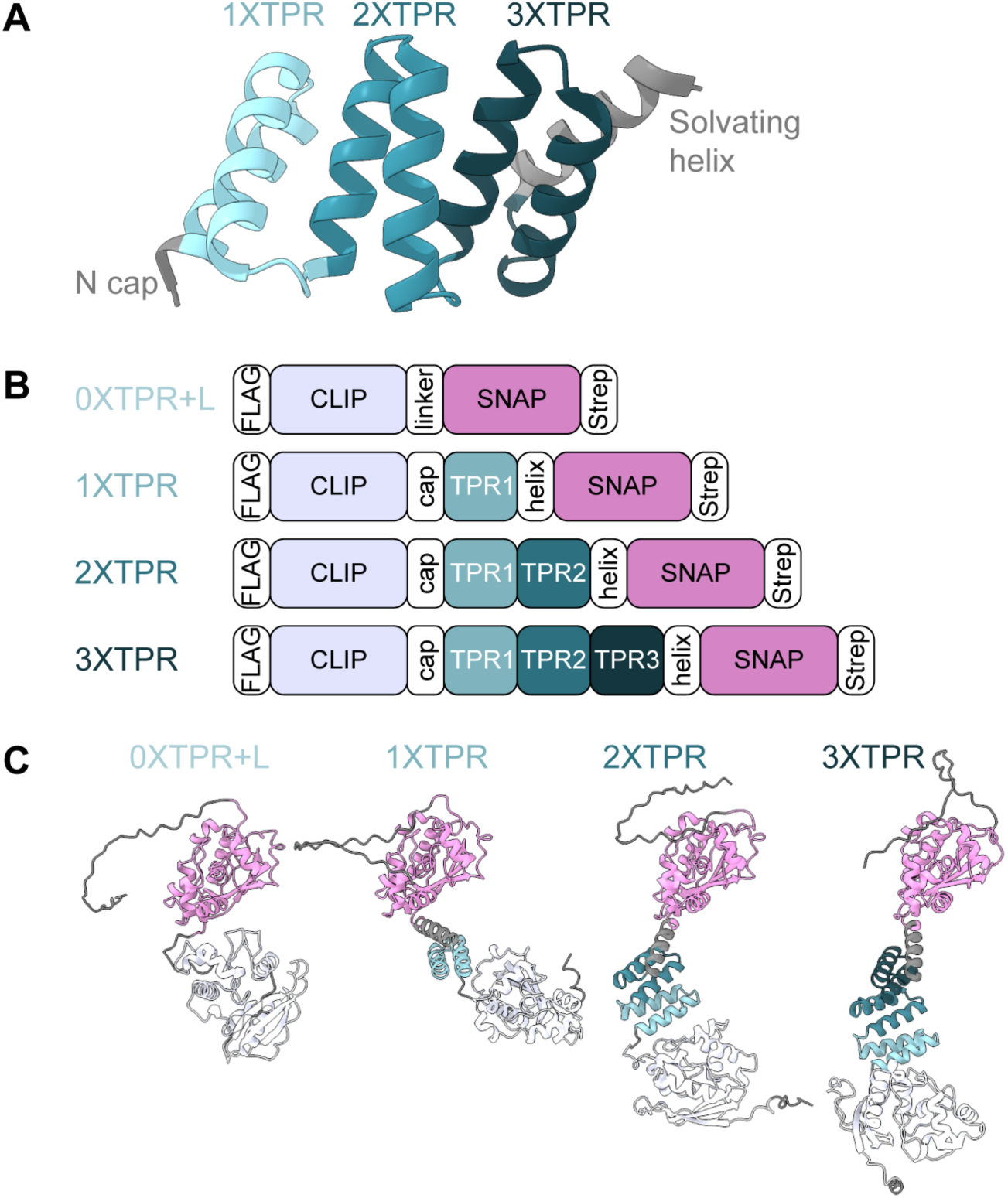
Design of protein standard ladder using self-labelling enzymes and tetratricopeptide repeat motifs. (A) Structure of TPR3 showing the three TPR motifs, the N cap and the solvating helix (PDB: 1na0, ^18^). (B) Schematic of four protein FRET standards making up the protein standard ladder; 0XTPR+L, 1XTPR, 2XTPR and 3XTPR. (C) AlphaFold prediction of the four protein FRET standards, domains are coloured as in the schematic above (B).

To provide a labelling system equally accessible *in vitro* and in cells, we used self-labelling enzymes. These bind covalently to ligands to which fluorescent organic dyes can be attached. The ligands can be cell permeable, meaning self-labelling enzymes can be labelled intracellularly and as purified protein. For single molecule work, these dyes have superior photophysical properties, compared to fluorescent proteins, with a much more uniform fluorescent signal ^19–21^. The variety of fluorescent ligands available for these tags also allows flexibility in the choice of FRET fluorophore pairs, depending on the restrictions of the instrumentation available.

The final FRET protein ladder constructs use the self labelling enzymes CLIP and SNAP flanking the repeatable TPR motif (Figure 1A, Figure 1B & Figure S1). The CLIP tag is a derivative of the SNAP-tag, with the same small size (19.4 kDa) maintaining the FRET dynamic range. The smallest construct (0XTPR+L) is a fusion of CLIP and SNAP connected by a short 12 amino acid linker (R-T-A-G-S-A-A-G-S-G-I-D). Increasing numbers of the idealised TPR motifs are placed in between the two tags to create a ladder (1XTPR-3XTPR); this increases the distance between the fluorophores, decreasing the FRET efficiency. An N-terminal FLAG tag and a C-terminal StrepTag were included for protein purification and immuno-assays.

The first FRET assay used to characterise the protein ladder was confocal single molecule FRET (smFRET). Here the absolute FRET efficiency, E, is calculated using alternating laser excitation (ALEX). A single molecule diffuses through the confocal volume at any one time, whilst lasers that excite the donor or the acceptor fluorophores alternate during the experiment resulting in bursts of photons from labelled molecules. Based on the excitation at any one time, emission photons can be split into three channels; donor-emission under donor excitation (DD), acceptor-emission under acceptor excitation (AA) and acceptor-emission under donor excitation (DA). FRET efficiency and stoichiometry are calculated, ranging from 0 to 1, from the number of photons in each of these channels. Calibration data provides *α*, *β*, *δ*, and *γ* correction factors to turn the apparent FRET efficiency into an absolute FRET efficiency.

The designed standards were codon optimised for mammalian expression, inserted into pcDNA3.1(+) and transfected into human 293 FT cells. We have previously shown that smFRET measurements can be taken directly from overexpressed labelled protein in diluted mammalian cell lysate ^8^. The four constructs were labelled with CLIP-Cell TMR-Star and SNAP-Cell 647- SiR using our previously optimised labelling protocol ^8^. Cells were lysed, centrifuged at 17, 000 x g for 10 minutes at 4 °C, and the supernatant diluted for single molecule concentrations (usually 1:100,000). Labelling was confirmed and non-specific labelling (which could result in an increase in background signal) was checked by ingel fluorescence (Figure S2A).

All four FRET standards resulted in a consistent smFRET population (Figure 2A to Figure 2D). Stoichiometry of labelling for all samples was around 0.5 showing equal labelling of the donor and the acceptor: 0XTPR+L 0.52 *±* 0.11, 1XTPR 0.53 *±* 0.12, 2XTPR 0.53 *±* 0.12 and 3xTPR 0.54 *±* 0.12 (mean *±* sd). This can also be seen by plotting the cumulative frequency of the stoichiometry, where the vast majority of molecules have a stoichiometry between 0.25 and 0.75 (Figure 2E). When analysing the FRET efficiency results, data is filtered with a stoichiometry between these values to select for dual labelled molecules.

**Figure 2:**
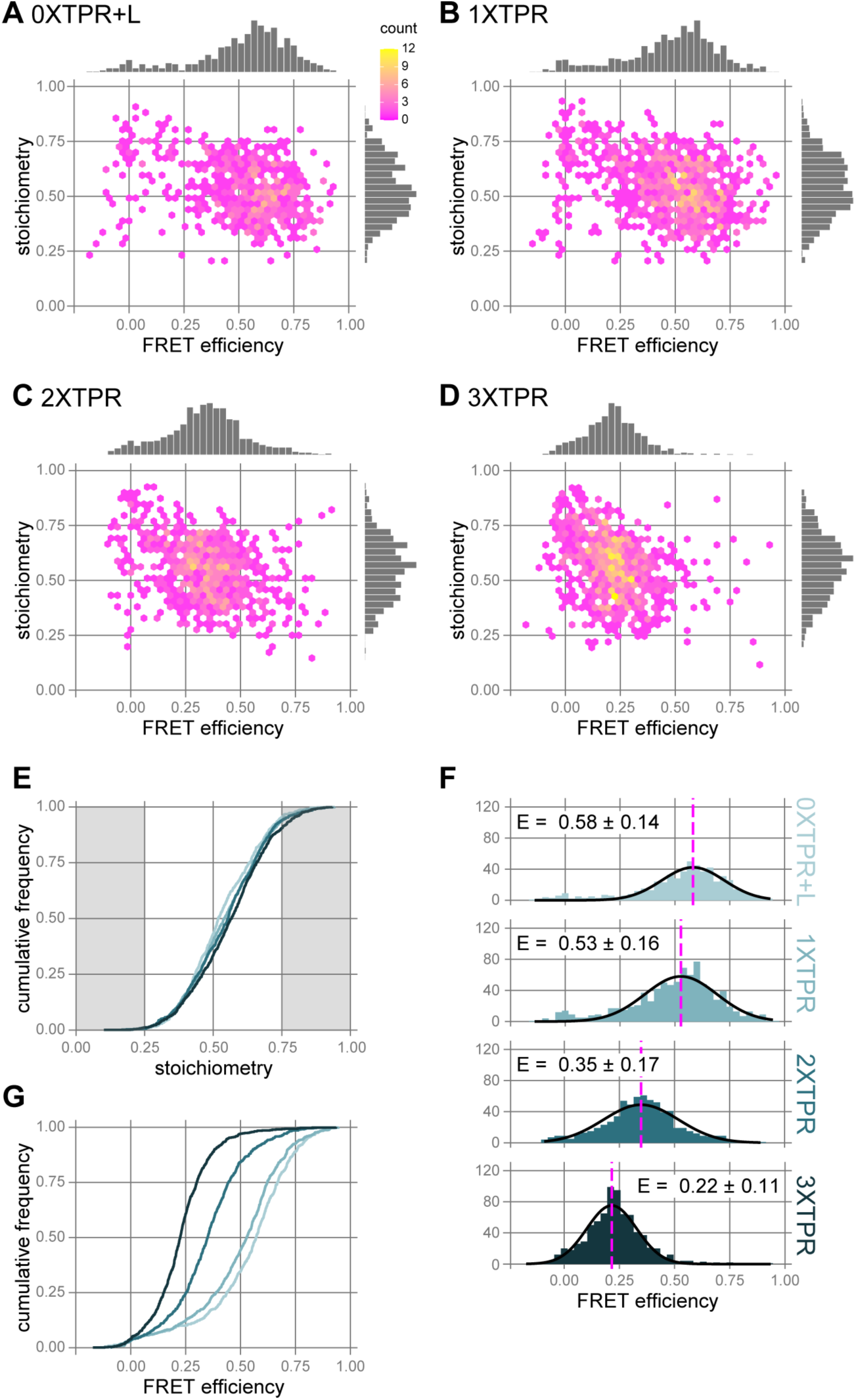
Samples from cell lysate result in a FRET ladder in smFRET. (A-D) Hexbin plots of FRET efficiency versus stoichiometry for overexpressed CLIP-SNAP fusion protein (0XTPR+L, A), CLIP-1XTPR-SNAP fusion protein (1XTPR, B), CLIP-2XTPR-SNAP fusion protein (2XTPR, C) and CLIP-3XTPR-SNAP fusion protein (3XTPR, D) from 293FT cell lysate. Data is combined from two to four biological replicates, n = 591, 917, 747 and 860 respectively. (E) Cumulative frequency plot for overexpressed 0XTPR+L, 1XTPR, 2XTPR and 3XTPR comparing stoichiometry from (A-D). Stoichiometry less than 0.25 and greater than 0.75 are highlighted in grey. (F) Histograms of FRET efficiency for overexpressed 0XTPR+L (n = 558), 1XTPR (n = 850), 2XTPR (n = 697) and 3XTPR (n = 768). Stoichiometry was filtered between 0.25 and 0.75 (G) Cumulative frequency plot comparing FRET efficiencies from (F) for overexpressed 0XTPR+L, 1XTPR, 2XTPR and 3XTPR. Pairwise Kolmogorov-Smirnov tests with Bonferroni correction resulted in p values of < 2.2e^−16^ for all comparisons apart from between 0XTPR+L and 1XTPR where p = 0.0036. Stoichiometry was filtered between 0.25 and 0.75.

Each FRET standard had a distinct FRET efficiency, and together the ladder spanned the FRET efficiency range (Figure 2F). As expected, the smallest construct of the CLIP-SNAP fusion protein (0XTPR+L) resulted in the highest FRET efficiency with mean *±* sd of 0.58 *±* 0.12. Then with increasing numbers of TPR repeat motifs, increasing the distance between the fluorophores, the FRET efficiency decreased: 1XTPR = 0.53 *±* 0.16; 2XTPR = 0.35 *±* 0.17; and 3XTPR resulting in the lowest FRET efficiency of 0.22 *±* 0.11 (mean *±* sd). This result can be seen more clearly in the plot of cumulative frequency where increasing the size of the FRET standard shifts the FRET efficiency to the left (Figure 2G). These results were consistent across replicates (Figure S2B & Figure S2C). However, the relative decrease in FRET efficiency from 0XTPR+L to 1XTPR was smaller than that of 1xTPR to 2XTPR and 2XTPR to 3XTPR.

In initial trials we also tested a fusion protein of the HaloTag and the SNAP-tag. HaloTag has a high labelling efficiency compared to the CLIP tag, minimising fluorophore labelling times and concentrations ^22^. 293FT cells were transfected and labelled with HaloTag TMR and SNAP-Cell 647-SiR. After analysis the mean FRET efficiency of HaloTag-SNAP was only 0.3 *±* 0.19, meaning this approach lacked the dynamic range needed to make a FRET standard ladder (Figure S2D). This is due to the relative positions of the bound fluorescent ligands and the larger size of the HaloTag (33 kDa) compared to the SNAP-tag (Figure S2E, Figure S2F).

### The FRET ladder has consistent FRET efficiency, regardless of expression system

In order for the FRET ladder to work for a diverse array of applications, E should be consistent across expression systems and labelling strategies. To increase the FRET efficiency of the smallest construct, we removed the flexible linker between the CLIP and SNAP tags to create a bigger difference in E when compared to 1XTPR. This construct (0xTPR), along with 1XTPR, 2XTPR and 3XTPR, was codon optimised for bacterial expression, inserted into pRESTA and transformed into BL21 E. coli cells. Protein expression was induced using IPTG and each construct purified by Strep-Tactin affinity chromatography (Figure S3A). To optimise efficient double labelling of purified protein, the shortest FRET standard construct, 0XTPR, was used. The fluorescent ligands CLIP-Cell TMR-Star and SNAP-Cell 647-SiR were added to the protein at differing protein:ligand molar ratios and left overnight at 4 °C. Labelling was measured by ingel fluorescence (Figure S3B). A protein:ligand molar ratio of 1:5 for both ligands was determined to be the most efficient and cost effective labelling strategy.

Each of the purified FRET standards, labelled with the two fluorescent ligands (Figure S3C), resulted in a consistent smFRET population (Figure 3A to Figure 3D). Again stoichiometry for each was around 0.5 (Figure 3E): 0XTPR = 0.52 *±* 0.12; 1XTPR = 0.51 *±* 0.12; 2XTPR = 0.50 *±* 0.12; and 3xTPR = 0.50 *±* 0.12 (mean *±* sd).

**Figure 3:**
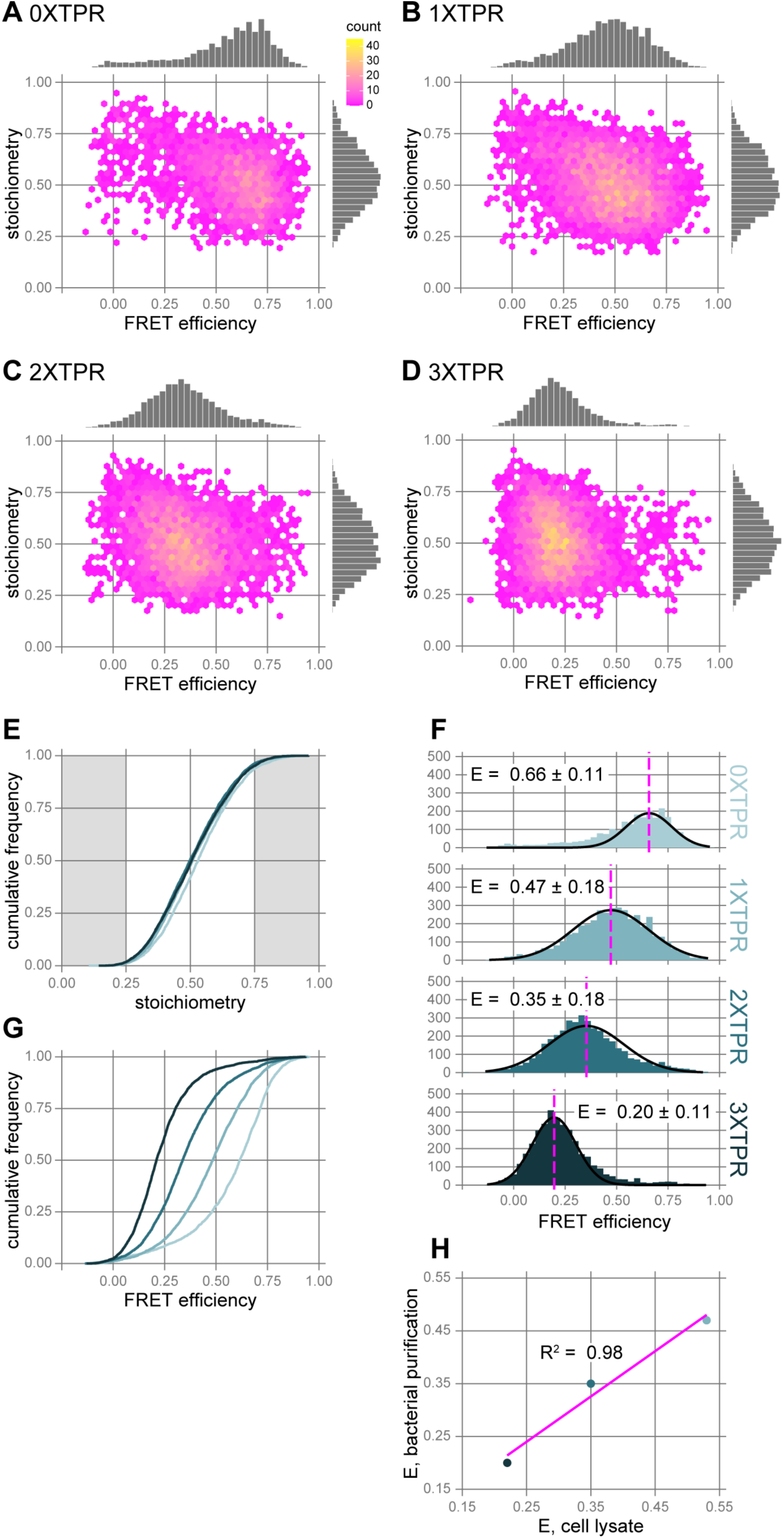
The FRET ladder is consistent for smFRET regardless of expression system. (A-D) Hexbin plots of FRET efficiency versus stoichiometry for purified CLIP-SNAP fusion protein (0XTPR, A), CLIP-1XTPR-SNAP fusion protein (1XTPR, B), CLIP-2XTPR-SNAP fusion protein (2XTPR, C) and CLIP-3XTPR-SNAP fusion protein (3XTPR, D) from E. coli. Data is combined from three to four biological replicates, n = 2613, 4482, 4102 and 4069 respectively. (E) Cumulative frequency plot for purified 0XTPR, 1XTPR, 2XTPR and 3XTPR comparing stoichiometry from (A-D). Stoichiometry less than 0.25 and greater than 0.75 are highlighted in grey. (F) Histograms of FRET efficiency for purified 0XTPR (n = 2422), 1XTPR (n = 4214), 2XTPR (n = 3880) and 3XTPR (n = 3799). Stoichiometry was filtered between 0.25 and 0.75. (G) Cumulative frequency plot comparing FRET efficiencies from (F) for purified 0XTPR, 1XTPR, 2XTPR and 3XTPR. p values calculated by pairwise Kolmogorov-Smirnov tests with Bonferroni correction were all < 2.2e^−16^ for all comparisons. Stoichiometry was filtered between 0.25 and 0.75. (H) Comparison between FRET efficiency calculated by smFRET from purified (3F) and overexpressed (2F) for 1XTPR, 2XTPR and 3XTPR. Linear model fit shown in magenta.

The FRET efficiency results were consistent with those of overexpressed protein from 293FT cell lysate, with the ladder spanning the FRET efficiency range (Figure 3F). The smallest construct of the CLIP-SNAP fusion protein (0XTPR) resulted in the highest FRET efficiency with mean *±* sd of 0.66 *±* 0.11 and increasing numbers of TPR repeat motifs giving rise to decreasing FRET efficiencies: 1XTPR = 0.53 *±* 0.16; 2XTPR = 0.35 *±* 0.17; and 3xTPR = 0.22 *±* 0.11 (mean *±* sd). Again this result can be seen more clearly in the plot of cumulative frequency where increasing the size of the FRET standard shifts the FRET efficiency to the left (Figure 3G). The results are consistent over biological replicates (Figure S3D & Figure S3E).

Having used the same protein constructs (1XTPR, 2XTPR and 3XTPR) and fluorophores for mammalian and bacterial smFRET experiments, we are able to directly compare FRET efficiencies across sample preparations using the same instrument (Figure 3H & Figure S3F). We applied a linear regression model which showed highly consistent results across the ladder (*R*^2^ = 0.98), despite very different sample preparation techniques. In general, the samples made from large scale purification from E. coli were more stable over the acquisition period than those from 293 FT cell lysate, allowing more single molecule events to be recorded overall. However, both preparations are suitable for data acquisition.

### The FRET ladder is compatible with non-canonical amino acid labelling through click chemistry

Self labelling enzymes are readily labelled intracellularly making them a tractable tool for single molecule work, but their size creates significant limitations. Tag size effectively subtracts from the dynamic range of FRET, as well as limiting the places where tags can be incorporated without impacting protein structure. To have the ability to translate between in vitro smFRET results that inform on structural dynamics and cell based assays, the ladder should be compatible with site specific labelling techniques. We tested non-canonical amino acid (ncAA) labelling via click chemistry using genetic code expansion, mutating the glutamine residue at position 149 in the CLIP tag to the amber stop codon TAG (0XTPR+L(Q149), Figure 4A). When expressed in cells this results in the premature truncation of the protein. However when cotransfected with an orthogonal aminoacyl-tRNA synthetase/tRNA pair (tR-NAPyl/NESPylRSAF) and an early release factor (eRF1-(E55D)), and with addition of the ncAA (TCO*A), then the ncAA is incorporated into the protein at the position of the amber stop codon ^23,24^. This ncAA can then be fluorescently labelled by copper-free click chemistry via a Strain-Promoted Inverse Electron-Demand Diels-Alder Reaction (SPIEDAC) through the strained alkyne group of the TCO*A reacting with the tetrazine group of the ligand ^25^.

**Figure 4:**
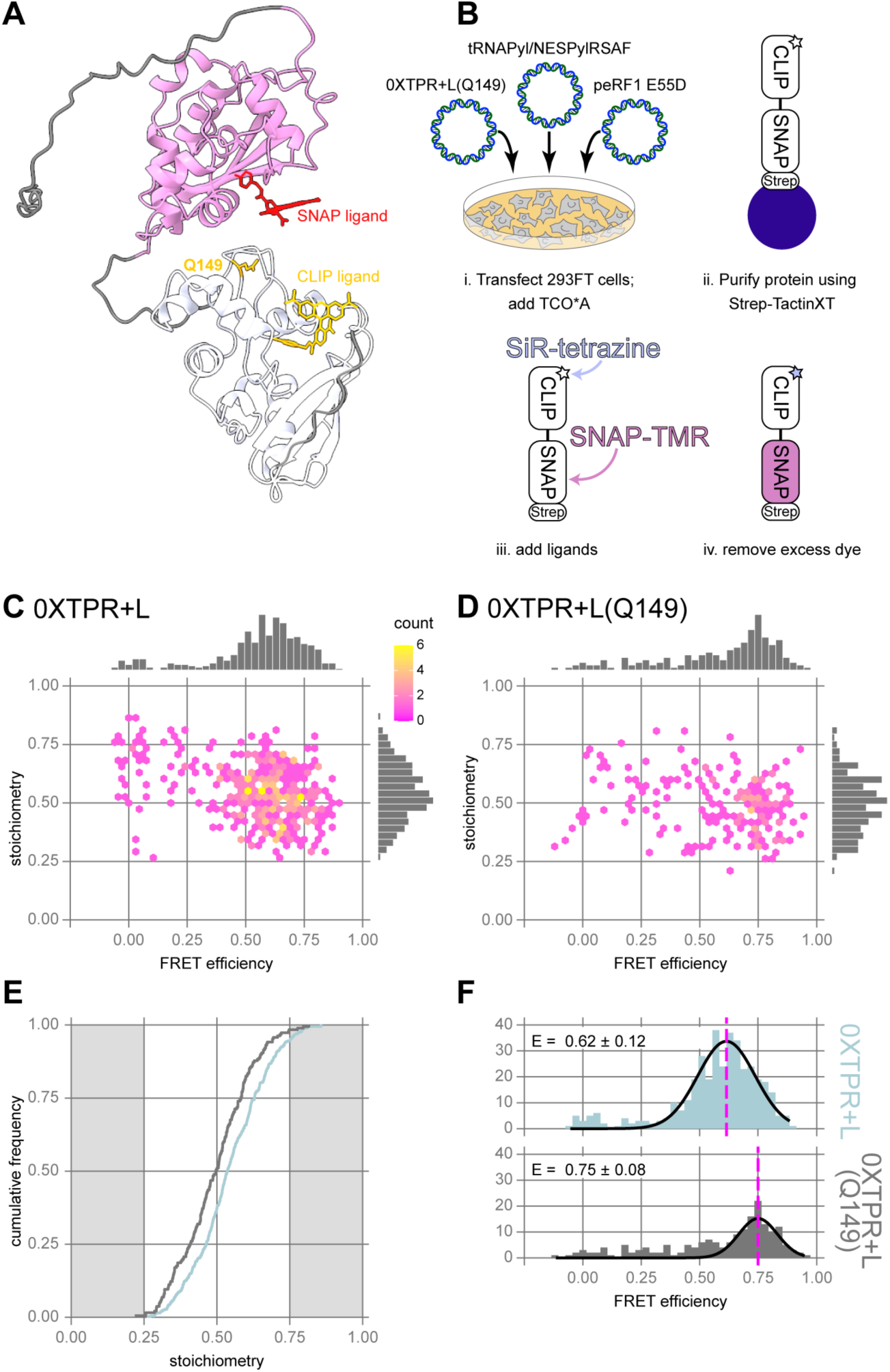
Non-cannonical labelling can be used with the FRET standard ladder. (A) AlphaFold prediction of the CLIP-SNAP fusion protein with a short flexible linker between the two tags (0XTPR+L). Domains are coloured as in schematic (1B) with Q149 highlighted. Ligands were placed based on PDB: 6y8p ^22^. (B) Steps for overexpression, small batch purification and double labelling of 0XTPR+L(Q149) with SNAP-Cell TMR Star and SiR-tetrazine. (C-D) Hexbin plots of FRET efficiency versus stoichiometry for purified CLIP-SNAP fusion protein (0XTPR+L, C) and CLIP(Q149)-SNAP fusion protein (0XTPR+L(Q149), D) from 293FT cells . Data from one biological replicate, n= 402 and 188 bursts respectively. (E) Cumulative frequency plot comparing the stoichiometry of 0XTPR+L (blue) and 0XTPR+L(Q149) (grey). Stoichiometry less than 0.25 and greater than 0.75 are highlighted in grey. (F) Histograms of FRET efficiency for purified 0XTPR+L (n = 380) and 0XTPR+L(Q149) (n = 184). Stoichiometry was filtered between 0.25 and 0.75.

0XTPR+L(Q149) was transfected into 293 FT cells, together with pcDNA3.1-tRNAPyl/NESPylRSAF and peRF1-E55D (Figure 4B). Cells were treated for eight hours with 2.5 mM of the ncAA (TCO*A) or DMSO as a control (Figure S4A). Complete read-through of protein was measured by labelling the C-terminal SNAP tag with SNAP-Cell TMR-Star and the cell lysate analysed by ingel fluorescence. This showed little read-through of the construct without the presence of TCO*A as expected, but good expression at the expected molecular weight for TCO*A treated cells. Addition of the TCO*A and TMR to untransfected cells resulted in no fluorescent ligand indicating no non-specific integration of the ncAA and fluorescent ligand.

We purified 0XTPR+L(Q149) and 0XTPR+L from transfected and TCO*A treated 293FT cells by small batch purification utilising the C-terminal StrepTag (Figure S4B). The fluorescent ligands SNAP-Cell TMR-Star and SiR-tetrazine were used to label the 0XTPR+L(Q149) protein, with the SiR-tetrazine labelling the ncAA by copper-free click chemistry (Figure 4B). The 0XTPR+L protein was labelled with CLIP-Cell TMR-Star and SNAP-Cell 647-SiR as previously (Figure S4C).

Both 0XTPR+L(Q149) and 0XTPR+L resulted in a high smFRET signal (Figure 4C & Figure 4D) with a mean stoichiometry of around 0.5: 0XTPR+L = 0.53 *±* 0.12; 0XTPR+L(Q149) = 0.49 *±* 0.11 (mean *±* sd). Based on the cumulative frequency plot of stoichiometry, apparent stoichiometry was different between 0XTPR+L(Q149) and 0XTPR+L, which we attribute to the different photophysical properties of the acceptor fluorophore (Figure 4E). 0XTPR+L(Q149) showed an increased FRET efficiency with a mean *±* sd of 0.75 *±* 0.08 compared to 0.62 *±* 0.12 for 0XTPR+L (Figure 4F). This is expected due to the different position of the donor fluorophore when using the non-cannonical labelling, positioning Q149 slightly closer to the SNAP tag in an elongated molecule (Figure 4A). Nonetheless, the FRET ladder can be successfully adapted to non-cannonical labelling approaches.

Strep-Tactin purification of 0XTPR+L from 293 FT cells provides an alternative sample preparation method to that used previously. Comparing the FRET efficiencies from Strep-Tactin purified molecules and those in diluted cell lysate shows highly reproducible FRET efficiency measurements; E, from cell lysate is 0.58 *±* 0.14 (mean *±* sd) whilst E of Strep-Tactin purified molecules is 0.62 *±* 0.12 (Figure S4D).

### Applying the FRET ladder to ensemble intracellular FRET assays

Our aim is to create a FRET ladder that allows the comparison of FRET results from single molecule *in vitro* work to ensemble measurements taken in cells. To understand the variability in intensity based FRET measurements in ensemble cellular assays, we used flow cytometry based-FRET. Flow cytometry is capable of assessing the fluorescence intensity of many thousands of labelled cells in only a few minutes. Thus, measuring FRET by flow cytometry allows for a much higher throughput assay. This has been used to screen protein-protein interactions ^26–28^ and protein conformation ^29^. However, these experiments mainly label the proteins of interest with fluorescent proteins and usually do not calculate FRET efficiencies. Instead either the percentage of FRET positive cells is measured, the ratio of acceptor to donor fluorescence is compared, or sensitised emission is calculated. 293 FT cells were transiently transfected with one of the four FRET standards or cotransfected with free CLIP and SNAP tags as a negative (no FRET) control. Cells were labelled as for smFRET assays with CLIP-Cell TMR-Star and SNAP-Cell 647-SiR. Single channel controls consisted of cells transfected with 0XTPR+L and labelled with only one of the fluorescent ligands resulting in donor only and acceptor only controls. Single living cells were selected by gating results based on the forward and side scatter (Figure S5A).

Doubly labelled cells were then gated from this population based on the donor and acceptor fluorescent channels (Figure 5A). The gate was set based on the single channel acceptor only control as the donor only control showed some bleed through of the donor emission into the acceptor channel. Doubly labelled cells were further gated to select FRET positive cells (Figure 5B); a triangular gate was created to exclude the no FRET control cells cotransfected with CLIP and SNAP tags. Gating resulted in over 97 % of doubly labelled cells being FRET positive in all four of the FRET constructs, compared to only 1 % for the cotransfected no FRET control cells (Figure 5C).

**Figure 5:**
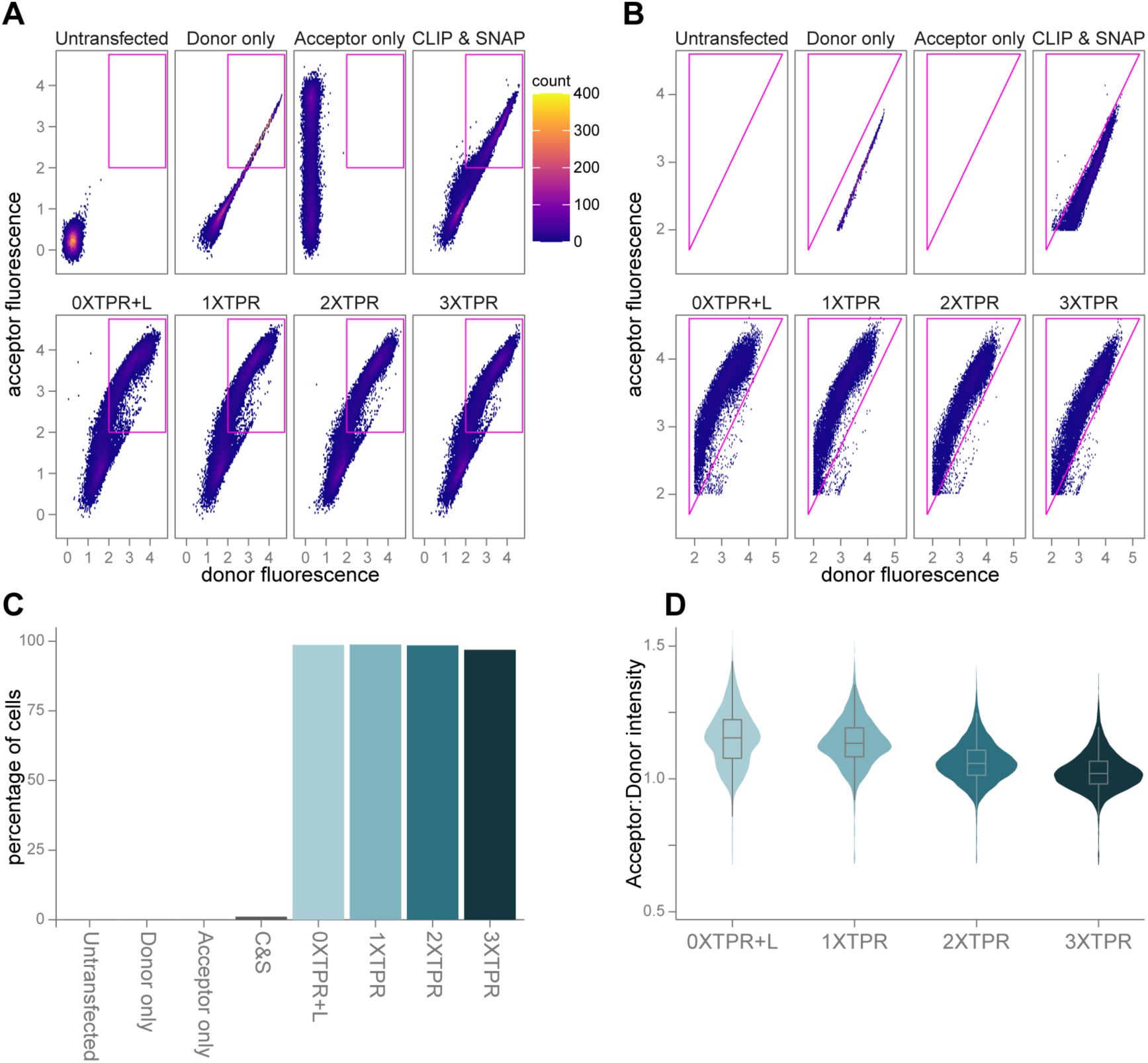
Using the FRET standards in flow cytometry based-FRET. (A) Hexbin plot of donor versus acceptor fluorescence of the single live gated cells for untransfected cells (n = 24,889), 0XTPR+L donor only (n = 29,456), 0XTPR+L acceptor only (n = 29,345), CLIP & SNAP cotransfection (n = 29,193), 0XTPR+L (n = 29,797), 1XTPR (n = 29,339), 2XTPR (n = 27,259) and 3XTPR (n = 27,525). Gate to select doubly labelled cells is shown in magenta. (B) Hexbin plot of donor versus acceptor fluorescence of the doubly labelled gated cells for untransfected cells (n = 0), 0XTPR+L donor only (n = 12,076), 0XTPR+L acceptor only (n = 0), CLIP & SNAP cotransfection (n = 13,354), 0XTPR+L (n = 15,431), 1XTPR (n = 15,615), 2XTPR (n = 13,504) and 3XTPR (n = 14,304). Gate to select FRET positive cells is shown in magenta. (C) Percentage of doubly labelled cells in the FRET positive gate for untransfected cells (0 %), 0XTPR+L donor only (0 %), 0XTPR+L acceptor only (0 %), CLIP & SNAP cotransfection (1 %), 0XTPR+L (99 %), 1XTPR (99 %), 2XTPR (98 %) and 3XTPR (97 %). (D) Ratio of acceptor:donor fluorescence intensity for 0XTPR+L, 1XTPR, 2XTPR and 3XTPR doubly labelled cells. All comparisons have a p value of <2e^−16^ (Kruskal-Wallis test then post-hoc pairwise Wilcoxon test with bonferroni correction).

We took a ratiometric approach to compare FRET efficiency along the ladder by looking at the ratio of acceptor to donor fluorescence intensity for each doubly labelled cell (Figure 5D). The ratio decreased across the FRET standard constructs as the distance between the fluorophores increased with the number of TPR units. The shortest construct 0XTPR+L resulted in the largest ratio of 1.15 *±* 0.11, followed by 1XTPR at 1.14 *±* 0.09, 2XTPR at 1.06 *±* 0.08 and the largest construct 3XTPR with the smallest ratio, 1.03 *±* 0.07 (mean *±* sd). All comparisons between constructs were significantly different using the Kruskal-Wallis test and post-hoc pairwise Wilcoxon tests with Bonferroni correction. This demonstrates that the FRET ladder can be used in flow cytometry based-FRET assays, both for a binary FRET/no FRET result, but also for quantitative measures of relative FRET across the dynamic range.

FRET impacts the fluorescence lifetime of both donor and acceptor fluorophores and so can be measured in Fluorescence Lifetime Imaging Microscopy (FLIM) experiments. Importantly FLIM can be used to determine sub-cellular distribution of complexes within cells ^30^, protein-protein interactions ^31^ and protein conformation ^32^. In this technique the fluorescence lifetime of the donor fluorophore is measured in the presence and absence of the acceptor fluorophore. If FRET is occurring between the pair of fluorophores this quenches the donor fluorescence and shortens the fluorescence lifetime. Measuring donor lifetime in the time domain by Time-Correlated Single Photon Counting (TCSPC), means lifetimes arent altered by fluorescence intensity or concentration.

COS-7 cells were transfected with one of the four FRET standards and labelled with CLIP-Surface 488 and SNAP-Cell TMR-Star post PFA fixation. As a negative (no FRET) control, cells were cotransfected with free CLIP and SNAP tags as before for the flow cytometry based-FRET experiments. Single channel controls consisted of CLIP and SNAP cotransfected cells labelled with either CLIP-Surface 488 (donor only) or SNAP-Cell TMR-Star (acceptor only). Successful labelling was confirmed prior to FLIM imaging by confocal microscopy of the same cells (Figure 6A and Figure S6A). This showed minimal bleed through of the donor fluorescence into the acceptor channel and minimal excitation of the acceptor fluorophore by the donor laser. Untransfected cells labelled with both fluorophores were used as a negative control to demonstrate the absence of non-specific fluorophore labelling and complete washout of unbound fluorescent ligands.

**Figure 6:**
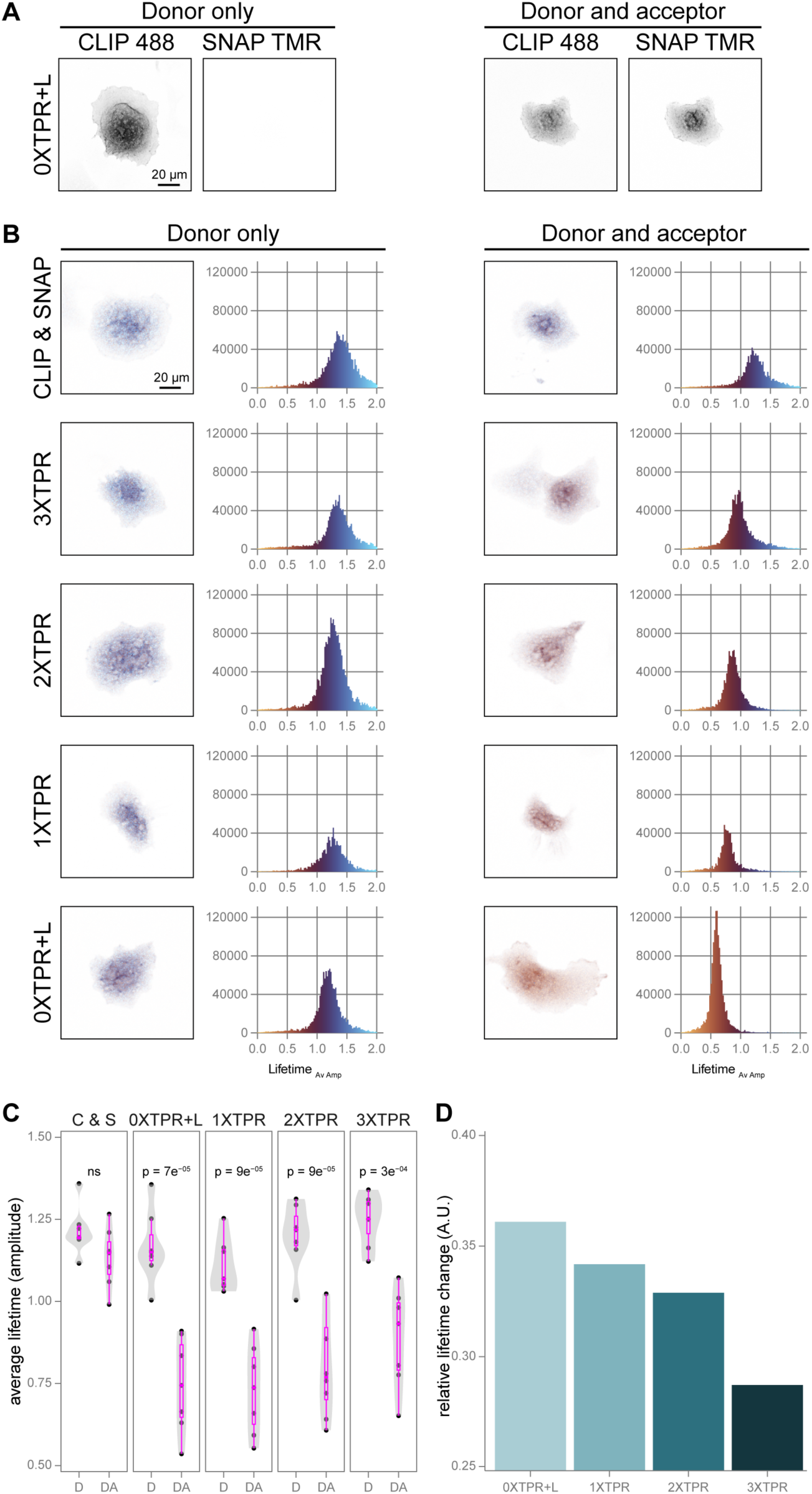
Using the FRET standards in flow cytometry based-FRET. (A) Representative confocal images of COS-7 cells transfected with the 0XTPR+L and labelled with only CLIP-Surface 488 (donor only, left) or both CLIP-Surface 488 and SNAP-Cell TMR-Star (donor and acceptor, right). (B) A representative cell coloured by average lifetime (amplitude) per pixel and the corresponding lifetime histogram for each FRET standard or a cotransfection of CLIP and SNAP. Cells were labelled only with CLIP-Surface 488 (donor only, left) or with SNAP-Cell TMR-Star as well (donor and acceptor, right). (C) Average lifetime (amplitude) of the four FRET standards; 0XTPR+L (n = 7 for D and DA), 1XTPR (n = 7 for D and DA), 2XTPR (D n = 7, DA n = 8), 3XTPR (n = 7 for D and DA) or a cotransfection of CLIP and SNAP (n = 7 for D and DA). For each construct, cells were labelled with just CLIP-Surface 488 (D, left) or also with SNAP-Cell TMR-Star (DA, right). p values were calculated by t-test. (D) Relative lifetime change for each of the four FRET standards calculated by 1-(mean lifetime DA/mean lifetime D).

The donor fluorescence lifetime was measured for each of the four FRET standards and the cotransfection of CLIP and SNAP, for the doubly labelled and donor only samples (Figure 6B). All four FRET standards show a significant decrease in donor lifetime when both fluorophores were present, compared to donor labelling alone (Figure 6C). The mean *±* sd for each FRET standard is reduced from: 1.17 *±* 0.11 to 0.74 *±* 0.14 for 0XTPR+L; 1.11 *±* 0.08 to 0.73 *±* 0.14 for 1XTPR; 1.20 *±* 0.10 to 0.80 *±* 0.16 for 2XTPR; and 1.25 *±* 0.08 to 0.89 *±* 0.15 for 3XTPR. In control experiments, there was no significant difference in donor lifetime for any of the FRET standard samples compared to the cotransfection of CLIP and SNAP when only the donor fluorophore was present (Figure S6B). There was also no significant difference in average donor lifetime for the negative control, cotransfection of CLIP and SNAP, between the donor only samples (mean *±* sd, 1.12 *±* 0.07) and when the acceptor fluorophore was also present (1.13 *±* 0.09).

To determine the difference between each of the FRET standards, we calculated the relative lifetime change between the donor only samples and those with both fluorophores (Figure 6D). Across the FRET standards this change was reduced as the distance between the fluorophores increased with the number of TPR units present: 0XTPR+L = 0.36; 1XTPR = 0.34; 2XTPR = 0.33; and 3XTPR = 0.29.

### The FRET ladder can correlate measurements across single molecule and ensemble assays

Using the same standardised FRET ladder across experiments opens up the possibility of correlating absolute measures of FRET efficiency in confocal smFRET assays, with relative changes observed within cells. To demonstrate the feasibility of this approach, we applied linear regression models to both the ratio of acceptor:donor intensity obtained from flow cytometry based-FRET (Figure 7A) and the relative lifetime change from FLIM-FRET (Figure 7B) with the confocal smFRET data from cell lysate assays. The models had *R*^2^ values of 0.98 and 0.90 respectively, demonstrating the utility of the ladder to correlate FRET quantification across acquisition systems, despite highly divergent acquisition techniques.

**Figure 7:**
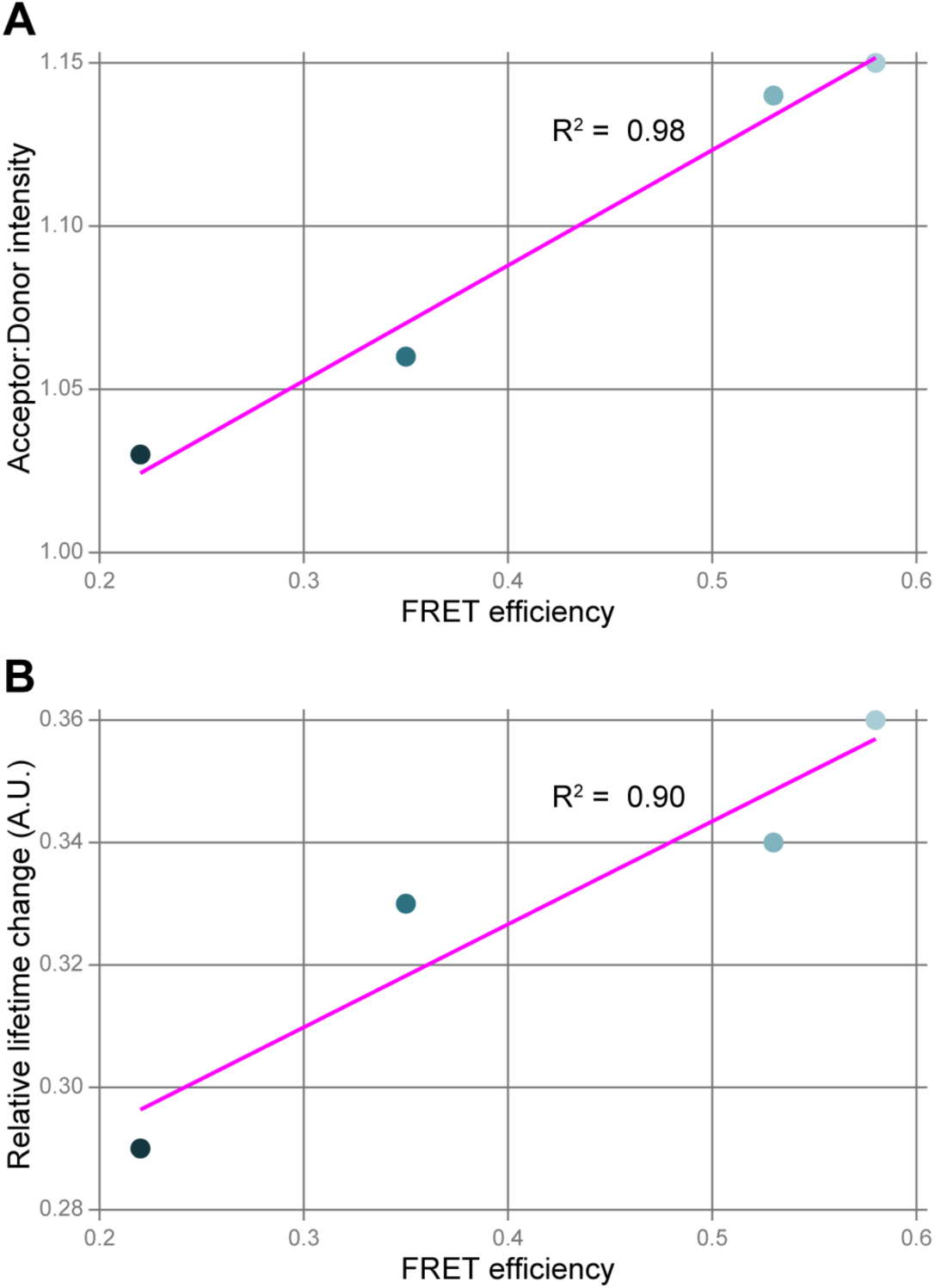
Using the FRET standards in flow cytometry based-FRET. (A) Comparison between FRET efficiency calculated by smFRET and the ratio of acceptor:donor intensity calculated by flow cytometry based-FRET for each of the four FRET standards; 0XTPR+L, 1XTPR, 2XTPR and 3XTPR. Linear model fit shown in magenta. (B) Comparison between FRET efficiency calculated by smFRET and the relative lifetime change calculated by FLIM-FRET for each of the four FRET standards. Linear model fit shown in magenta.

## DISCUSSION

We have developed a protein standard ladder that can be used in a diverse set of FRET based assays. The four standards, which only differ by the number of TPR repeat motifs between the CLIP and SNAP tags (0XTPR, 1XTPR, 2XTPR and 3XTPR), result in a range of FRET efficiencies from around 0.2 to 0.7. The design of the standards means that multiple sample preparation methods can be used; here we have demonstrated measurements taken directly in cell lysate (Figure 2), large/small scale protein purification (Figure 3 and Figure 4 respectively) and measurement in intact cells, with (Figure 6) and without fixation (Figure 5). The self labelling enzymes CLIP and SNAP allow for both in-cell and *in vitro* labelling, whilst engineered TPR motifs are highly stable and unrelated to any human proteins. The use of the CLIP and SNAP tags allows the choice of the donor and acceptor fluorophores for FRET to be changed depending on the equipment at hand. In this work we focussed on the cell permeable ligands: CLIP-Cell TMR-Star, SNAP-Cell TMR-Star and SNAP-Cell 647-SiR but, brighter non-cell-permeable fluorescent ligands could have been used for the purified protein samples. We have shown that the standards can be adapted by a simple mutagenesis PCR to other site specific labelling techniques; in this case non-canonical amino acid labelling using genetic code expansion. The protein ladder gave highly comparable quantitative FRET measurements in all three FRET assays tested; confocal smFRET, flow cytometry based-FRET and FLIM-FRET.

### Adapting the protein FRET standards to other labelling strategies

When designing the protein FRET standards we found that the size of the self labelling enzyme affected the effective dynamic range of FRET efficiencies that are available (Figure S2D). Consequently we chose the CLIP and SNAP tags (19.4 kDa), rather than the larger HaloTag, even though the labelling efficiency of the CLIP tag has more variability and slower kinetics ^8,22^. The smaller size of SNAP/CLIP allowed the standards to span most of the FRET efficiency range, however an even smaller self-labelling enzyme, BromoCatch ^33^, is now available which could further improve the high FRET efficiency limit beyond 0.7 achieved here.

We have tested two different labelling approaches in this work, self labelling enzymes and non-canonical amino acid labelling. However, other strategies could be used with only minor adaptations to the FRET protein ladder. Maleimide labelling is commonly used for studies using purified proteins, through labelling of surface available cysteine residues ^34^. Both the CLIP and SNAP tags naturally contain three cysteine residues, but the engineered TPR motifs do not. Modified standards could be made by removing the CLIP and SNAP tags and adding cysteine residues to the N- or C- terminus of the different sized TPR motifs. Similarly the 11-residue ybbR peptide ^35^ could be added to provide orthogonality to the labelling of the N- and C- terminii, facilitating site specific linkage of donor and acceptor fluorophores. Although this labelling approach is only applicable to purified protein samples and couldnt be used in cell based assays, it may still prove useful for correlating across instruments dedicated to in-solution versus surface immobilised FRET measurements ^13^. Similarly, the SNAP and CLIP tags could be replaced by intrinsically fluorescent proteins to quantify the dy-namic range of new cellular biosensors that are developed ^36,37^.

As a fluorescence lifetime measurement, FLIM-FRET is independent of fluorescence intensity ^38,39^. However fluorescence lifetime is altered by the local environment ^40–42^. Here, we demonstrate variability in the lifetime measurements due to differences in local environments of the donor fluorophore following over-expression of the protein ladder throughout the cell (Figure 6C). Consequently, we would recommend addition of sub-cellular localisation motifs, either directly or by fusion to a marker protein, to allow the protein ladder to be used within a targeted local environment relevant to the biological question at hand. This would ensure any change in lifetime was due to FRET, with the protein ladder standardising measurements within the local environment.

### Protein ladder applications

The advent of nanoscopy techniques demanded characterisation of reference structures or rulers to estimate the spatial resolution of these super-resolution microscopy methods. Endogenous nuclear pore complexes (NPCs) emerged as resolution standards for calibration and quality control within cells ^43,44^, in parallel with the engineered DNA origami structures that have become in-valuable tools for demonstrating microscope performance for *in vitro* samples ^45,46^. These have been supplemented with homo-oligomeric protein standards to support accurate protein counting ^47^ and protein reference structures to validate resolution ^48^. Critically, these structures have also provided readily accessible benchmarks to support new users and the development of imaging and analysis pipelines facilitating uptake of techniques. They allow experimentalists to master new methods without having to troubleshoot the biology and the method at the same time. Those wishing to apply advanced FRET measurements to protein samples have not previously had access to the same facility.

There are three specific use cases where we are already applying the FRET protein ladder to support our own work. First, in smFRET studies, calculating *γ* and *β* correction factors requires a range of measured FRET efficiencies, preferably with samples for both low FRET and high FRET ^10^. The protein ladder provides a multipoint calibration tool to supply these measurements that can be integrated with the same sample preparation techniques as the experimental molecules with the same fluorophores. This avoids relying on the experimental molecules to provide the dynamic range needed, which may not be possible for many studies. Second, the most basic but perhaps the most impactful, onboarding new users with advanced biophysics analysis techniques. For example, during confocal smFRET experiments when arriving at the instrument for data collection, the abstract nature of the raw data can make it difficult to immediately determine if sample preparation, labelling, laser and instrument as a whole are all working and consistent across replicates. Likewise, if a single component of this workflow is faulty, successfully isolating the problem without pos=itive control molecules that have been prepared by the same techniques is almost impossible. This is in stark contrast to light microscopy of cells, where the user has immediate access to qualitative measures of sample integrity and labelling before having to rely on the detection modules of the instrument in question. Finally, as the protein ladder behaves as a standard curve to link measurements that would otherwise be unrelated to each other, it facilitates the interpolation of FRET based signals between very different acquisition platforms to understand protein conformations not only in vitro, but within their cellular context.

In summary, we have established a modular, biologically compatible ‘protein ladder’ that serves as a universal calibration curve for FRET. By bridging the gap between *in vitro* biophysics and live-cell physiology, this tool provides a robust framework for the interpretation of molecular distances across diverse FRET modalities. Consequently, this platform facilitates the integration of high-resolution structural states resolved by smFRET with their functional behaviour in the native cellular environment.

## REAGENTS AND TOOLS TABLE

Key reagents and their sources are listed in Table 1.

**Table 1:** Reagents and tools Key reagents used in this study and their source.

| Reagent/Resource | Reference or Source | Identifier or Catalogue Number |
| --- | --- | --- |
| **Experimental models** |  |  |
| 293FT cell line | ThermoFisher Scientific | R70007 |
| COS-7 cell line | ECACC | 87021302 |
| **Recombinant DNA** |  |  |
| pCDNA3.1-FLAG-CLIP-SNAP-StrepTag | this study | Addgene #248177 |
| pCDNA3.1-FLAG-CLIP-1XTPR-SNAP-StrepTag | this study | Addgene #248178 |
| pCDNA3.1-FLAG-CLIP-2XTPR-SNAP-StrepTag | this study | Addgene #248179 |
| pCDNA3.1-FLAG-CLIP-3XTPR-SNAP-StrepTag | this study | Addgene #248180 |
| pRESTA-FLAG-CLIP-SNAP-StrepTag | this study | Addgene #248181 |
| pRESTA-FLAG-CLIP-1XTPR-SNAP-StrepTag | this study | Addgene #248182 |
| pRESTA-FLAG-CLIP-2XTPR-SNAP-StrepTag | this study | Addgene #248183 |
| pRESTA-FLAG-CLIP-3XTPR-SNAP-StrepTag | this study | Addgene #248184 |
| pCDNA3.1-FLAG-CLIP(Q149TAG)-SNAP-StrepTag | this study | Addgene #248185 |
| pCDNA3.1-CLIP-HisTag-FLAG | 8 | Addgene #200787 |
| pCDNA3.1-SNAP-HisTag-FLAG | 8 | Addgene #200788 |
| pCDNA3.1-FLAG-HaloTag-SNAP-StrepTag | this study | Addgene #248186 |
| pcDNA3.1-tRNAPyl/NESPyIRSAF | Kind gift from Edward Lemke | 23 |
| peRF1 E55D | Kind gift from Jason Chin | 24 |
| **Antibodies** |  |  |
| Anti-SNAP-tag antibody | New England Biolabs | P9310S |
| HRP conjugated -Rabbit IgG | Strattech | 211-032-171 |
| **Chemicals, Enzymes and other reagents** |  |  |
| Biotin | IBA/Merck | 2-1016-005/B4501-10G |
| 4-12 % Bis-Tris gel | ThermoFisher Scientific | NP0335BOX |
| B-PER Complete Bacterial Protein Extraction Reagent | ThermoFisher Scientific | 15255743 |
| BSA | Sigma-Aldrich | AO281 |
| CLIP-Cell TMR-Star | New England Biolabs | S9219S |
| CLIP-Surface 488 | New England Biolabs | S9232S |
| Dulbeccos Modified Eagle Medium (DMEM) - high glucose | Sigma-Aldrich | D5796 |
| Fetal Bovine Serum (FBS) | Sigma-Aldrich | F7524 |
| Formaldehyde 16% | TAAB Laboratory and Microscopy | F017 |
| Geneticin | Appleton Woods | CSR552 |
| GlutaMAX, 100X | ThermoFisher Scientific | 11574466 |
| HaloTag TMR Ligand | Promega | G8251 |
| Halt protease inhibitor cocktail | ThermoFisher Scientific | 11824111 |
| 1 M HEPES | ThermoFisher Scientific | 15630106 |
| IPTG | Merck | I5502-10G |
| LipoD293 DNA In Vitro Transfection Reagent | TEBU-Bio | 189SL100668-1ml |
| MagStrep Strep-TactinXT beads | ThermoFisher (IBA) | 25090010 |
| MEM Non-Essential Amino Acid Solution, 100X | ThermoFisher Scientific | 12084947 |
| MOPS running buffer (20X) | Fisher | 11589156 |
| NuPAGE LDS Sample Buffer (4X) | ThermoFisher Scientific | NP0007 |
| NuPAGE Sample Reducing Agent (10X) | ThermoFisher Scientific | NP0009 |
| Penicillin Streptomycin Solution, 50X | Sigma-Aldrich | P0781 |
| ProLong Gold antifade reagent | ThermoFisher Scientific | P36961 |
| Quick Coomassie Stain | Protein Ark | GEN-QC-STAIN-1L |
| SiR-Tetrazine | TEBU-Bio | SC008-1 |
| SNAP-Cell 647-SiR | New England Biolabs | S9102S |
| SNAP-Cell TMR-Star | New England Biolabs | S9105S |
| Sodium Pyruvate | Gibco | 11360070 |
| StrepTactinXT 4Flow high capacity 1 ml column | ThermoFisher Scientific | 17285794 |
| TCO<sup>a</sup>, [(E)-cyclooct-2-en-1-yl]oxycarbonyl-L-lysine | SiChem | SC-8008 |
| Trans-blot turbo mini 0.2 μm nitrocellulose transfer packs | Biorad | 1704158 |
| Trypsin-EDTA 0.5 % Solution, 10X | Santa Cruz Biotechnology | sc-363354 |
| Turbo DNase | Invitrogen | AM2238 |
| Zeba dye and biotin removal spin column | ThermoFisher Scientific | A44297 |
| **Software** |  |  |
| AlphaFold 3 | https://alphafoldserver.com/welcome |  |
| Fiji/ImageJ | https://imagej.net/software/fiji/ |  |
| FRETbursts python package | https://fretbursts.readthedocs.io/ |  |
| Jupyter Notebook | https://jupyter.org/ |  |
| RStudio | https://cran.r-project.org/ |  |
| SymPhoTime 64 software v2.8 | https://www.picoquant.com/products/category/software/symphotime-64-fluorescence-lifetime-imaging-and-correlation-software |  |
| **Other** |  |  |
| AKTA Pure 25 L system | Cytiva |  |
| BD FACS Melody | BD Biosciences |  |
| Custom built smfBox | 49 |  |
| EI-FLEX | Exciting Instruments |  |
| G:box gel imager | Syngene |  |
| Odyssey Fc Imager | LI-COR |  |
| LSM980 AiryScan 2 confocal microscope | Zeiss |  |

## MATERIALS AND METHODS

### Recombinant DNA expression plasmids

There are two sets of FRET standards; either codon optimised for mammalian or bacterial expression. The mammalian expression plasmids were all synthesised by GeneArt (Invitrogen by ThermoFisher Scientific) resulting in the four constructs: pCDNA3.1-FLAG-CLIP-3XTPR-SNAPf-StrepTag, pCDNA3.1-FLAG-CLIP-2XTPR-SNAP-StrepTag, pCDNA3.1-FLAG-CLIP-1XTPR-SNAP-StrepTag, pCDNA3.1-FLAG-CLIP-SNAP-StrepTag. This construct, 0XTPR+L, has a short 12 amino acid linker R-T-A-G-S-A-A-G-S-G-I-D between the CLIP and SNAP tags. Site-directed mutagenesis PCR of this construct was performed to introduce a TAG stop codon within the CLIP tag replacing Glutamine 149. This resulted in the pCDNA3.1-FLAG-CLIP(Q149TAG)-SNAP-StrepTag construct (0XTPR+L(Q149)). Single CLIP and SNAP only plasmids (pCDNA3.1-CLIP-HisTag-FLAG and pCDNA3.1-SNAP-HisTag-FLAG) have been previously described ^8^ and were used as no FRET controls. The pCDNA3.1-FLAG-HaloTag-SNAP-StrepTag construct has the same short linker between the two tags and was also synthesised by GeneArt. pRESTA-FLAG-CLIP-3XTPR-SNAPf-StrepTag (codon optimised for bacterial expression) was synthesised by GeneArt. Deletion mutagenesis PCR was used to create the three shorter constructs; pRESTA-FLAG-CLIP-2XTPR-SNAP-StrepTag, pRESTA-FLAG-CLIP-1XTPR-SNAP-StrepTag, pRESTA-FLAG-CLIP-SNAP-StrepTag (0XTPR). All plasmids above are available from Addgene (see Materials availability). pcDNA3.1-tRNAPyl/NESPylRSAF was a gift from Edward Lemke ^23^ and peRF1 E55D was a gift from Jason Chin ^24^.

### Bacterial expression and large scale protein purification

Constructs codon optimised for bacterial expression were transformed into BL21 E.coli. Single colonies were inoculated into 50 ml of LB culture, with 50 *µ*g/ml carbenicillin and 35 *µ*g/ml chloramphenicol and grown overnight at 37 °C with shaking at 220 RPM. Overnight cultures were used to inoculate 1 litre of Terrific Broth (TB; 12 g/L Tryptone, 24 g/L Yeast extract, 0.4 % (vol/vol) Glycerol, 17 mM KH_2_PO_4_ and 72 mM K_2_HPO_4_), supplemented with the same antibiotic concentrations. Cultures were grown at 37 °C with shaking at 220 RPM until the optical density reached 0.8. Cultures were then cooled to 18 °C before protein expression was induced with 0.2 mM IPTG (isopropyl *β*-D-1-thiogalactopyranoside); cultures were shaken at 220 RPM overnight at room temperature. To harvest the cells, the cultures were pelleted by centrifugation at 4000 x g for 15 minutes at 4 °C. The cell pellet was resuspended in 10 ml of 2xPBS + 1 mM DTT and centrifuged at 4000 x g for 20 minutes at 4 °C. The final pellet was flash frozen in liquid nitrogen and stored at −80 °C until needed.

One induced bacterial cell pellet was thawed and resuspended in B-PER Complete Bacterial Protein Extraction Reagent (5 ml per g of pellet) and Halt protease inhibitor cocktail (50 *µ*l per g of pellet). The cell suspension was incubated on a roller at room temperature for 20 minutes before clearing by centrifugation at 23,000 x g for 30 minutes at 4 °C. Affinity chromatography was carried out using an AKTA Pure 25 L system with sample application and elution taking place at a flow rate of 0.5 ml/min, with all other steps at 1 ml/min. Briefly, the supernatant was filtered, using a 0.20 *µ*m syringe filter, and loaded onto a StrepTactinXT 4Flow high capacity 1 ml column, preequilibrated with 5 column volumes (CVs) of wash buffer (100 mM Tris pH 8, 150 mM NaCl, 1 mM EDTA pH 8). The column was then washed with 10 CVs of wash buffer before a gradient elution up to a maximum of 50 mM biotin (100 mM Tris pH 8, 150 mM NaCl, 1 mM EDTA pH 8, 10 % glycerol, 50 mM biotin). 2 ml fractions from the elution peak were collected, aliquoted into single use aliquots and flash frozen in liquid nitrogen. Samples from each stage were analysed by SDS-polyacrylamide gel elec-trophoresis and Quick Coomassie Stain.

### Cell culture and transfection

293FT cells (Invitrogen) were cultured in Dulbeccos Modified Eagle Medium supplemented with 10 % (vol/vol) Fetal Bovine Serum, 2 mM GlutaMAX, 500 *µ*g/ml Geneticin, 1 mM Sodium Pyruvate, 1X MEM Non-Essential Amino Acid Solution, 500 U Penicillin and 500 *µ*g Streptomycin. COS-7 cells were cultured in Dulbeccos Modified Eagle Medium supplemented with 10 % (vol/vol) Fetal Bovine Serum, 500 U Penicillin and 500 *µ*g Streptomycin. All cells were grown at 37 °C with 5 % CO_2_ and checked regularly for mycoplasma contamination. Every 2-3 days when 80 % confluent cells were passaged by trypsinization. For smFRET, FACS-FRET and FLIM-FRET experiments, cells were plated at 50 % confluency in a 6-well plate. 18 hours later, cells were transfected using the LipoD293 DNA *in vitro* Transfection Reagent. 1 *µ*g of DNA was added to 50 *µ*l of DMEM and 3 *µ*l of transfection reagent was added to another 50 *µ*l of DMEM. The diluted transfection reagent solution was added dropwise to the diluted DNA solution. This mixture was left to in-cubate for 10 minutes at room temperature. Media was removed from the cells leaving only 1 ml and the DNA transfection mixture was added dropwise to the well.

### Genetic code expansion

[(E)-cyclooct-2-en-1-yl]oxycarbonyl)-L-lysine (TCO*A) was prepared to a concentration of 100 mM in 0.2M NaOH with 15 % DMSO, according to manufacturers instructions. This was diluted to 25 mM in 1 M HEPES and was added immediately after transfections to a final concentration of 2.5 mM. To test the read-through of protein TCO*A, 300,000 cells per well of 293FT cells were plated in a 6-well plate and after 24 hours transfected with LipoD293. Per well, 400 ng of both the 0XTPR+L(Q149) and the pcDNA3.1-tRNAPyl/NESPylRS constructs as well as 250 ng of peRF1-E55D (ratio of 1:1:0.6) was diluted in 50 *µ*l of DMEM. 3 *µ*l of LipoD293 was diluted in another 50 *µ*l of DMEM before combining with the diluted DNA. The mixture was incubated for 10 minutes at room temperature before adding to the cells. TCO*A was then added to cells and left to incubate for eight hours before the media was exchanged to fresh media. As a control DMSO was also diluted at a 1:4 ratio in 1 M HEPES. The following day SNAP-Cell TMR-Star was added to the cells at a final concentration of 0.5 *µ*M and left to incubate for two hours. The cells were then washed with media three times before incubation for two hours in 3 ml of fresh media. After the incubation the cells were washed in PBS, followed by a 30 second wash in Trypsin. The cells were incubated for 4 minutes at 37 °C, before harvesting in 1 ml DPBS and pelleting by centrifugation at 400 x g for 5 minutes at room temperature. The supernatant was discarded and 1 *µ*l of Turbo DNase was added to the pellet, before addition of 50 *µ*l of RIPA buffer (50 mM Tris HCl pH 7.3, 100mM NaCl, 2 mM MgCl, 1 % Triton X-100, 0.1 % Sodium deox-cholate, 0.1 % SDS) supplemented with 1 mM Dithiothreitol (DTT), 1 X Halt protease inhibitor cocktail, and 10X Turbo DNase buffer to a final concentration of 1X . Cells were incubated at 37 °C for 20 minutes, before centrifugation at 20,000 x g for 10 minutes at 4 °C. The supernatant was collected and 4X sample buffer and 10X reducing buffer was added to a concentration of 1X. Samples were then boiled at 70 °C for 15 minutes before analysis by in-gel fluorescence.

### Small batch protein purification

293FT cells were plated in 15 cm dishes, and transfected with LipoD293 after 24 hours. Per 15 cm dish, 12.5 *µ*g of DNA was di-luted in 625 *µ*l of DMEM and 37.5 *µ*l of LipoD293 was diluted in a further 625 *µ*l of DMEM. For amber suppression system exper-iments, cells were transfected with 4.75 *µ*g of both the 0XTPR+L(Q149) and the pcDNA3.1-tRNAPyl/NESPylRSAF constructs as well as with 3 *µ*g of peRF1-E55D (ratio of 1:1:0.6) to allow efficient incorporation of the non-canonical amino acid. The diluted DNA and LipoD293 were combined and incubated for 10 minutes before adding to the cells. For the smFRET experiments, two 15 cm dishes were transfected with 0XTPR+L and four with 0XTPR+L(Q149). TCO*A was added immediately after transfections to a final concentration of 2.5 mM and left on the cells for 48 hours. Cells were then harvested by washing the cells in PBS, followed by a 30 second wash in Trypsin. The cells were incubated for 4 minutes at 37 °C, before harvesting in 10 ml DPBS and pelleting by centrifugation at 400 x g for 5 minutes at room temperature. The supernatant was discarded and the pellet resuspended in 1 ml of PBS in a low-bind microcentrifuge tube before further centrifugation at 2500 x g at 4 °C for 10 minutes. The supernatant was removed and cell lysis was performed in NET buffer (50 mM Tris-Cl pH 7.3, 150 mM NaCl, 0.1 mM EDTA, 2 mM MgCl_2_, 2 mM CaCl_2_, 2 mM ZnCl_2_, supplemented with 0.1 % NP40, Halt Protease inhibitor cocktail and 10 mM sodium butyrate). 2 Units of Turbo DNase and 10 x Turbo DNase buffer were added directly to the resuspension and incubated at 37 °C for 30 minutes, until less viscous. Cells were centrifuged at 20,000 x g for 20 minutes at 4 °C. 200 *µ*l MagStrep Strep-TactinXT beads were washed three times in 200 *µ*l of Buffer W (100 mM Tris-HCl, 150 mM NaCl, 10 mM EDTA pH 8). The cell lysate was added to the washed beads and incubated with rotation for 2 hours at 4 °C. After incubation, the beads were washed 3 times in Buffer W before elution with 1 X BXT buffer (100 mM Tris-HCl, 150 mM NaCl, 2 mM EDTA, 100 mM biotin pH 8 supplemented with 5 % glycerol). Beads were incubated for 10 minutes at room temperature, vortexing 3 times. The supernatant was collected and flash frozen in liquid nitrogen before storage at −70 °C. Concentration of samples was determined by BCA assay. 4X sample buffer and 10X reducing buffer were added to samples from each stage to a concentration of 1X. Samples were analysed by SDS-PAGE and western blotting.

### Fluorescently labelling purified protein

To optimise the protein to ligand molar ratio for labelling, CLIP-Cell TMR-Star and SNAP-Cell 647-SiR were added to large scale purified 0XTPR at different ratios (1:0.5, 1:1, 1:2.5, 1:5, 1:12.5 and 1:25) keeping the concentration of protein constant at 0.5 *µ*M. Samples were left overnight at 4 °C before addition of 3X sample buffer (150 mM Tris pH 6.8, 6 % SDS, 0.3 M DTT, 0.3 % Bromophenol and 30 % glycerol) to a final concentration of 1X. These samples were then boiled at 100 °C for five minutes and analysed by in-gel fluorescence. A ratio of 1:5 was then used for all future labelling of large scale purified protein. For small batch purified protein, a higher ratio of 1:12.5 was used for both SNAP-Cell TMR-Star and SiR-Tetrazine. This was due to the lower concentration of 0XTPR+L(Q149) obtained. After overnight labelling at 4 °C, a Zeba dye and biotin removal spin column was used to remove any unbound dye. Briefly, the column was spun at 1000 x g for 2 minutes at 4 °C to remove the storage buffer before the labelled protein was slowly added to the centre of the resin. The column was then centrifuged again at 1000 x g for 2 minutes at 4 °C. The flow-through was collected in a low-bind microcentrifuge tube and used for smFRET assays.

### In-cell labelling with cell permeable ligands

Labelling conditions for CLIP-Cell TMR-Star and SNAP-Cell 647-SiR were optimised previously ^8^ and these conditions are used for the in-cell labelling in this publication. Briefly, CLIP-Cell TMR-Star was added to the 293FT cells six hours after transfection at a final concentration of 0.5 *µ*M and left to incubate at 37 °C with 5 % CO_2_ for 15 hours on the cells. SNAP-Cell 647-SiR was added the next day at the same final concentration and left to incubate for two hours. HaloTag TMR was added at a final concentration of 25 nM and left to incubate for 15 minutes. Cells were then washed twice with fresh media and incubated in 3 ml of media for two hours to remove any unbound dye. Cells were washed for the final time with 1 ml of PBS before incubating with 500 *µ*l of 0.5 % trypsin for 5 minutes. The trypsin was neutralised by addition of 1 ml of cell culture media. Cells were pelleted by centrifugation at 500 x g for 4 minutes at 4 °C. The pellet was resuspended in 1 ml of PBS before a further centrifugation step at 500 x g for 4 minutes at 4 °C. Cells were resuspended in 50 *µ*l of lysis buffer (40 mM HEPES pH 7.5, 1 mM EDTA pH 8, 120 mM NaCl, 0.05 % Triton X-100, 1 *µ*g/ml Aprotinin, 10 *µ*g/ml Leupeptin, 1 *µ*g/ml Pepstatin A, 10 *µ*g/ml TAME) and incubated on ice for 10 minutes followed by centrifugation at 17, 000 x g for 10 minutes at 4 °C. The resultant supernatant was used for smFRET assays.

### smFRET

Labelled purified protein or supernatant from cell lysis was diluted in smFRET assay buffer (40 mM Tris pH 8, 5 mM NaCl, 1 mM DTT, 0.1 mg/ml photobleached BSA) to obtain single molecule resolution. For lysate samples this was usually 1:100,000 and for purified protein 1:1×10^18^. Samples were analysed using a custom built smFRET confocal microscope, the smfBox ^49^. This consists of a laser, emission filter and avalanche photodiode for both the donor (80 mW 515 nm and 571/72) and the acceptor (100 mW 638 nm 678/41). The donor laser was used at 65 % and the acceptor at 12 % with these lasers alternating on and off over the duration of the experiment (either 15 or 30 minutes). Data was first analysed using a custom built Jupyter notebook and the FRET-Bursts package ^50^ to eliminate background and for burst selection (see data and code availability). Correction factors were calculated for each labelling condition. For the C(Q149)-S and HaloTag-SNAP data sets a range of FRET efficiencies was not measured for these fluorophore pairs, so gamma and beta were estimated as 1. The threshold for burst selection was set at 20 photons in both the DD+DA and AA channels and 20 photons above background for all constructs except HaloTag-SNAP where they were set at 20 and 10 respectively. Selected bursts were then analysed using a custom built R notebook for further filtering (see data and code availability). Any data sets with a burst rate over 1 burst per second were removed as being too concentrated and unlikely to be of single molecule resolution. For background elimination, filters were set at above: 16.5 photons for the donor excitation (DD+DA), 3 for donor emission (DD) and 6 for acceptor excitation and emission (AA). To eliminate noise, two gaussians were predicted when estimating the FRET efficiency for each construct unless the data was clean enough that only one gaussian was needed. The pairwise Kolmogorov-Smirnov tests were performed with Bonferroni correction to compare multiple cumulative frequencies. The labelling of all samples was checked after the experiment using in-gel fluorescence. When comparing the FRET efficiencies against other FRET measurements results, a linear regression model was fitted in R Studio and the unadjusted R-squared value recorded.

### FACS-FRET

Labelled transfected cells were harvested by trypsinisation and resuspended in 1 ml of PBS as described above. The cell suspension was then analysed using a BD FACS Melody flow cytometer using the 561 nm laser. CLIP-Cell TMR-Star emission was detected through the PE 582/15 filter and SNAP-Cell 647-SiR through the PE-Cy5 697/58 filter. After transformation the raw data was then gated by forward scatter and side scatter area to select single live cells. Further gating by donor and acceptor fluorescence intensity was done to select doubled labelled cells and then FRET positive cells. This was done using a custom R notebook (see data and code availability) and the flowcore ^51^, flowWorkspace ^52^, ggcyto ^53^ and ncdfFlow ^54^ packages.

### FLIM-FRET

COS-7 cells were plated in 6-well plates containing 13 mm acid washed glass coverslips and transfected as described above. 48 hours after transfection, cells were washed with PBS before fixation by paraformaldehyde incubation for 10 minutes at room temperature. Coverslips were then washed three times in PBS before incubation in permeabilisation solution (PBS with 0.1 % triton) for 5 minutes at room temperature. After five washes in PBS, coverslips were incubated overnight at 4 °C in 0.5 *µ*M of CLIP-Surface 488 and SNAP-Cell TMR-Star diluted in block solution (PBS with 5 % FBS, 1 % BSA). The five washes in PBS were repeated and the coverslips mounted onto clear glass slides using ProLong Gold antifade reagent and then sealed with nail varnish. All images were captured using the Zeiss LSM980 AiryScan 2 confocal microscope using the Zen Blue software v3.7. Confocal images were imaged with excitation at 488 nm for CLIP-Surface 488 and 561 nm for SNAP-Cell TMR-Star. The pinhole was set to 1 Airy unit resulting in 4.1 *µ*m and 4.9 *µ*m optical sections for each laser respectively. The frame size was 2048 x 2048 pixels and line average of eight. The brightness and contrast was set for each channel using ImageJ/Fiji ^55^ and the images prepared for publication using Adobe Illustrator. FLIM measurements were recorded using the TCSPC system (PicoQuant), the Zen Blue PicoQuant application and the SymPhoTime 64 software v2.8 (PicoQuant). The system has two PMA Hybrid 40 detectors (PicoQuant) with a 560 nm beam splitter and two emission filters (520/35 and 600/50 nm). The 485 nm laser at 40 MHz and 70 % intensity was used to measure the donor (CLIP-Surface 488) fluorescence lifetime. Laser power was altered depending on the level of expression/labelling of the chosen cell (typically 70-100 %) and the pinhole set to 2 Airy units resulting in a 8.2 *µ*m optical section. The FLIM measurements were taken using the following settings in T3 TCSPC mode: frame size of 256 x 256, pixel size of 829 nm, time resolution of 10 ps and a pixel dwell time of 32.77 *µ*s. Data was taken until the brightest pixel had 1000 photons (usually 5-10 minutes). The average fluorescence lifetime for each field of view was calculated in the SymPhoTime 64 software after fitting of the fluorescence decay curve with 3 parameters keeping the background parameters constant and the background of the decay set to zero. This data was extracted as an .ome.tiff file and processed for image presentation using ImageJ/Fiji ^55^ and a custom R notebook (see data and code availability). Packages used were: EBImage ^56^, scico ^57^, ggpubr ^58^ and purrr ^59^.

### In-gel fluorescence

To determine the protein labelling efficiency, samples were separated by 10 % SDS polyacrylamide resolving gels at 150 V for approximately one and a half hours. Gels were analysed using the Odyssey Fc Imager with CLIP-Cell TMR-Star/SNAP-Cell TMR-Star seen in the 600 channel and SNAP-Cell 647-SiR/SiR-Tetrazine in the 700 channel.

### Western blotting

Samples were prepared for western blotting by the addition of NuPAGE LDS sample buffer and NuPAGE sample reducing agent before boiling at 70 °C for 10 minutes. The samples were then loaded on 4-12 % Bis-Tris gel with MOPS running buffer (Fisher). Gel was transferred onto a membrane using trans-blot turbo mini 0.2 *µ*m nitrocellulose transfer packs (Biorad) before blocking with 2 % BSA in Tris-buffered saline-Tween. Membranes were probed with the anti-SNAP-tag primary antibody (1:2000) followed by HRP-conjugated anti-Rabbit secondary. Western blots were imaged using G:box (Syngene) gel imager. The brightness and contrast of images were adjusted using ImageJ/Fiji ^55^.

### Protein Structure Prediction

Three-dimensional protein structures of 3XTPR, 2XTPR, 1XTPR and 0XTPR+L were predicted using the AlphaFold 3 model ^60^ hosted on the Google AlphaFold server.

### Resource availability

#### Lead contact

Materials, data and code are available as indicated below. Any additional information or resource requests should be directed to the lead contacts, Evelyn Smith or Alison Twelvetrees.

#### Materials availability

All plasmids generated in this study are deposited with Addgene (addgene.org/browse/article/28262289/).

#### Data and code availability

All data and code used to make figures are deposited in ORDA, the University of Sheffield data repository, powered by Figshare (doi.org/10.15131/shef.data.c.8329780).

## ACKNOWLEDGMENTS

We thank Mahmoud A. S. Abdelhamid and Elliot M. Steele for their help with the smFRET experiments as well as Sophie E. Fountain for her help calculating the correction factors. We would also like to thank the Flow Cytometry Facility and the Wolfson Light Microscopy Facility at the University of Sheffield.

This work was supported through the following funding: E.R.S. was supported by a studentship from the White Rose BBSRC Doctoral Training Partnership (grant number: 2109768); E.R.S. also received a University of Sheffield Postgraduate Research Student Publication Scholarship; L. H-C received a Wellcome Trust Biomedical Vacation Scholarship; A.E.T. and D.A.B. are Sir Henry Dale Fellows, funded by the Wellcome Trust and the Royal Society (grant numbers: 220192/Z/20/Z and 213501/Z/18/Z). FLIM was performed at the Wolfson Light Microscopy Facility, using the Zeiss LSM980 Airyscan 2 Confocal microscope (MRC grant MR/X012077/1). D.A.B., A.E.T., T.D.C. and K.G. were supported by a BBSRC Pioneer Award (grant number: BB/Y513453/1).

This research was funded in whole, or in part, by the Wellcome Trust. For the purpose of Open Access, the author has applied a CC BY public copyright licence to any Author Accepted Manuscript version arising from this submission.

## AUTHOR CONTRIBUTIONS

Conceptualization: E.R.S, K.L.G., T.D.C., D.A.B., A.E.T.; Data curation: E.R.S., A.E.T.; Formal analysis: E.R.S., A.E.T.; Funding acquisition: E.R.S, T.D.C., D.A.B., A.E.T.; Investigation: E.R.S., K.L.G.; Methodology: E.R.S, K.L.G., L.H-C., T.D.C., D.A.B., A.E.T.; Project administration: A.E.T.; Supervision: T.D.C., D.A.B., A.E.T.; Resources: T.D.C., D.A.B., A.E.T.; Validation: E.R.S, K.L.G., T.D.C., D.A.B., A.E.T.; Visualisation: E.R.S.; Writing - original draft: E.R.S.; Writing - review and editing: all authors.

## AUTHOR COMPETING INTERESTS

T.D.C. is the founder and CEO of Exciting Instruments (EI), a company that develops and sells instrumentation for single-molecule fluorescence experiments, including smFRET. A.E.T. and E. R. S. previously held a EPSRC IAA grant in collaboration with EI.

## Notes

https://doi.org/10.15131/shef.data.c.8329780

https://www.addgene.org/Alison_Twelvetrees

### Summary of Updates

The title for Supplementary Figure 1 was incorrect and has now been changed. The table of resources now includes the final Addgene codes for the generated plasmids.

