## Supplementary Figures for "A universal protein ladder for standardisation of diverse FRET assays"

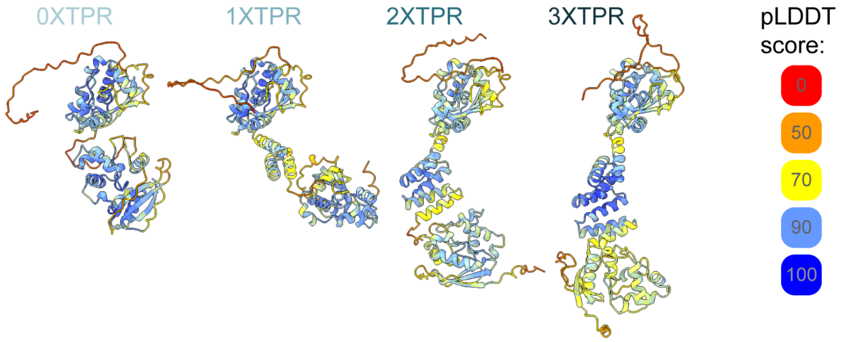


**Figure S1: AlphaFold confidence scores for the FRET protein ladder**
The same AlphaFold prediction shown in Figure 1C, coloured by pLDDT score.

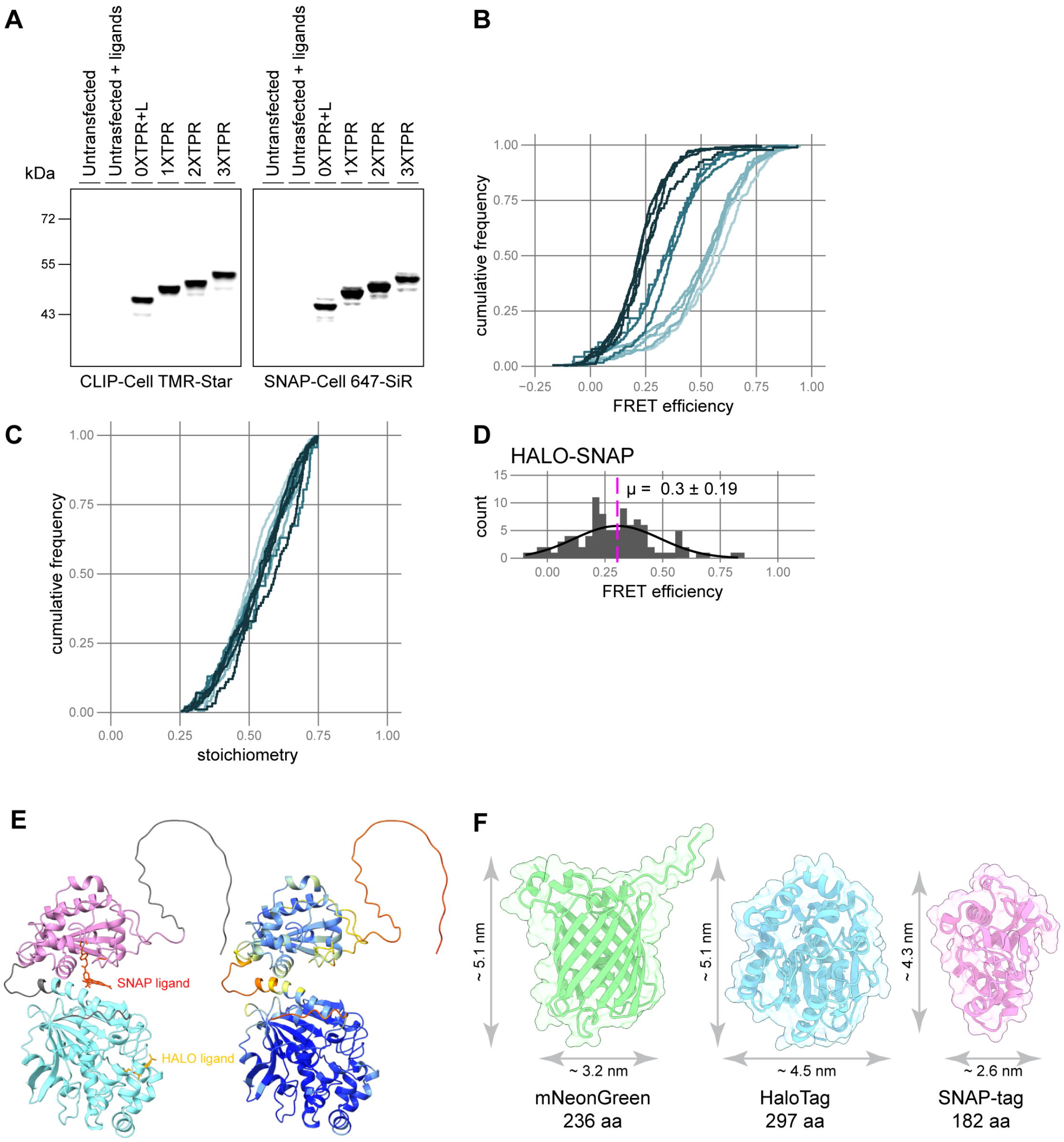


**Figure S2: smFRET results from cell lysate are consistent across biological replicates.**
(A) In-gel fluorescence of cell lysate from 293FT cells transfected with 0XTPR+L, 1XTPR, 2XTPR or 3XTPR. Cells were labelled with CLIP-Cell TMR-Star (left) and SNAP-Cell 647-SiR (right). Both untransfected controls, with and without addition of the two ligands, show no fluorescence.
(B, C) Cumulative frequency plot of FRET efficiency (B) and stoichiometry (C) for individual biological replicates of 0XTPR+L, 1XTPR, 2XTPR and 3XTPR. Stoichiometry was filtered between 0.25 and 0.75.
(D) FRET efficiency histogram of HALO-SNAP (n = 97 bursts) with no gating by stoichiometry.
(E) AlphaFold prediction of HALO-SNAP fusion protein; coloured by domain (left) and pLDDT score (right). PDB: 5y2y and 6y8p were used to place the HALO and SNAP ligands respectively [22,61].
(F) AlphaFold prediction of mNeonGreen, HaloTag and SNAP-tag showing the size difference between the three proteins.

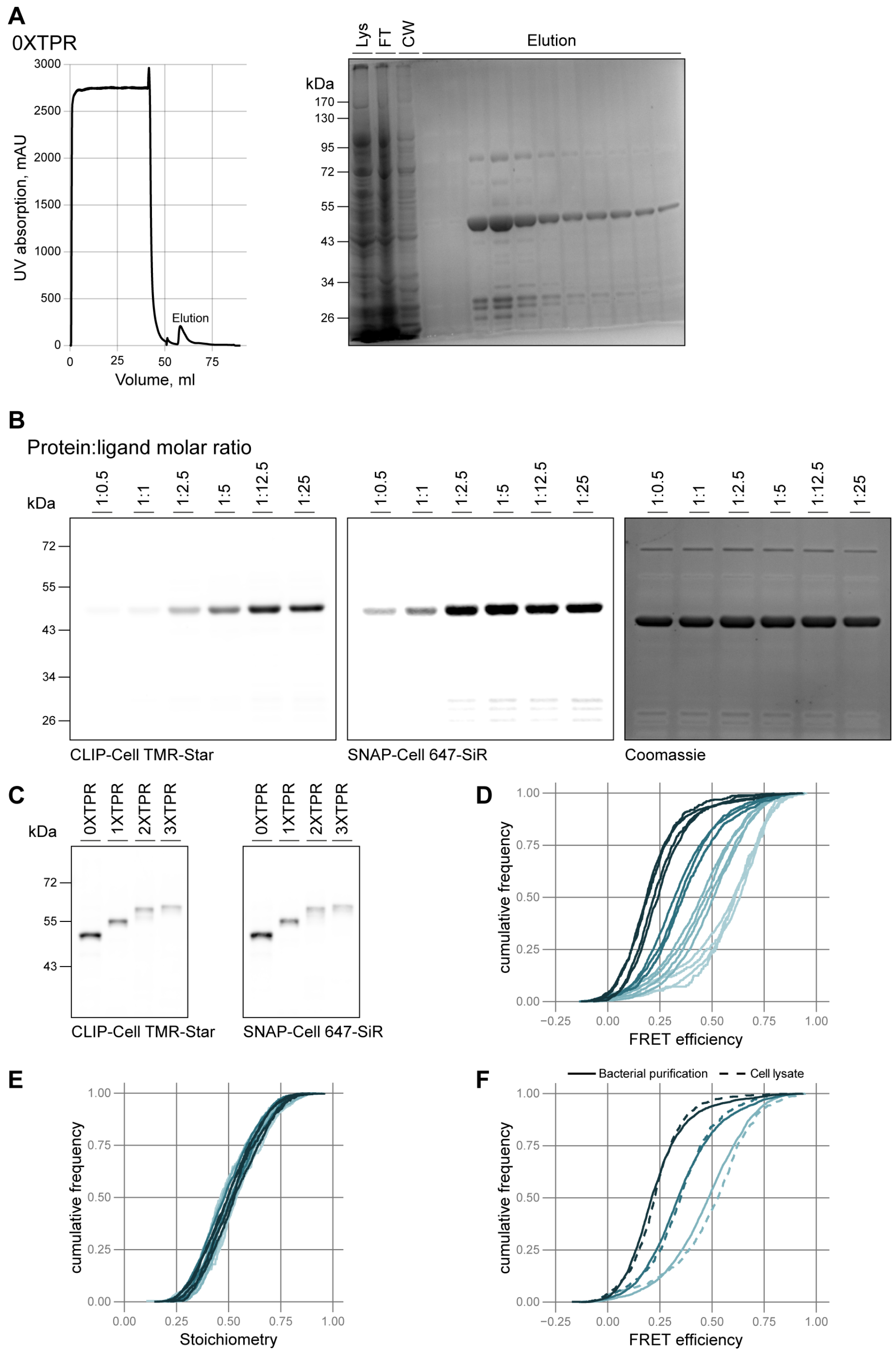


**Figure S3: Large scale purification from E.coli of the protein ladder results in consistent smFRET results.**
(A) Large scale purification of 0XTPR which is representative of that of 1xTPR, 2xTPR and 3XTPR. UV absorption during the protein purification performed on the AKTA Pure 25 L system (left) shows a peak during the elution phase. The purified protein can be seen just above 43 kDa on the Coomassie stained SDS-page gel (right). Samples: Lys, lysate; FT, flow through; CW, column wash; elution fractions.
(B) In-gel fluorescence of purified 0XTPR protein incubated with CLIP-Cell TMR-Star (left) and SNAP-Cell 647-SiR (middle) at the indicated protein:ligand molar ratios. The Coomassie stained gel is shown on the right.
(C) In-gel fluorescence of purified 0XTPR, 1XTPR, 2XTPR and 3XTPR labelled with CLIP-Cell TMR-Star (left) and SNAP-Cell 647-SiR (right) at a 1:5 protein:ligand molar ratio overnight.
(D-E) Cumulative frequency plot of FRET efficiency (D) and stoichiometry (E) for individual biological replicates of purified 0XTPR, 1XTPR, 2XTPR and 3XTPR. Stoichiometry was filtered between 0.25 and 0.75.
(F) Cumulative frequency plot comparing FRET efficiencies from purified (3F) and overexpressed (2F) for 1XTPR, 2XTPR and 3XTPR. Stoichiometry was filtered between 0.25 and 0.75.

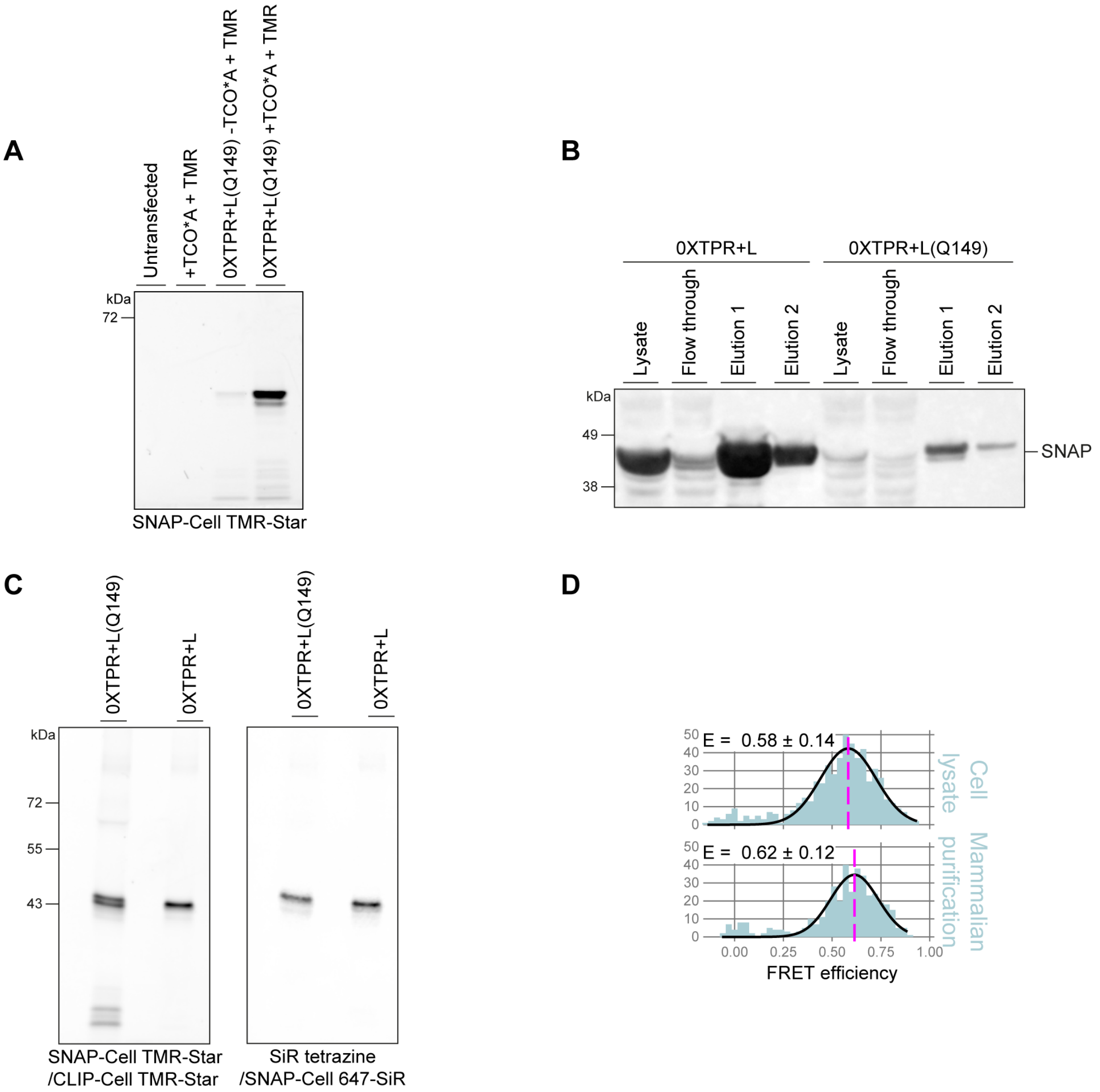


**Figure S4: Purification and labelling using non-canonical amino acid labelling through click chemistry.**

(A) In-gel fluorescence of cell lysate from 293FT cells looking at the translation read-through of 0XTPR+L(Q149) in the absence of the non-canonical amino acid (TCO*A). Cells were either left untransfected or transfected with the 0XTPR+L(Q149) construct and were labelled with SNAP-Cell TMR-Star.

(B) Small batch purification of 0XTPR+L and 0XTPR+L(Q149). 293FT cells were transfected with either of the two constructs and the overexpressed protein was purified using MagStrep Strep-TactinXT beads and biotin elution. Samples were separated by SDS-PAGE and analysed by western blot using the rabbit anti-SNAP-tag antibody. A much smaller amount of the 0XTPR+L(Q149) protein was expressed and therefore purified.

(C) In-gel fluorescence of purified 0XTPR+L(Q149), labelled with SNAP-Cell TMR-Star and SiR-tetrazine, and 0XTPR+L labelled with CLIP-Cell TMR-Star and SNAP-Cell 647-SiR.

(D) Histograms of FRET efficiency for overexpressed 0XTPR+L from 293FT cell lysate (2F) and purified 0XTPR+L from 293FT cells (4F). Stoichiometry was filtered between 0.25 and 0.75.

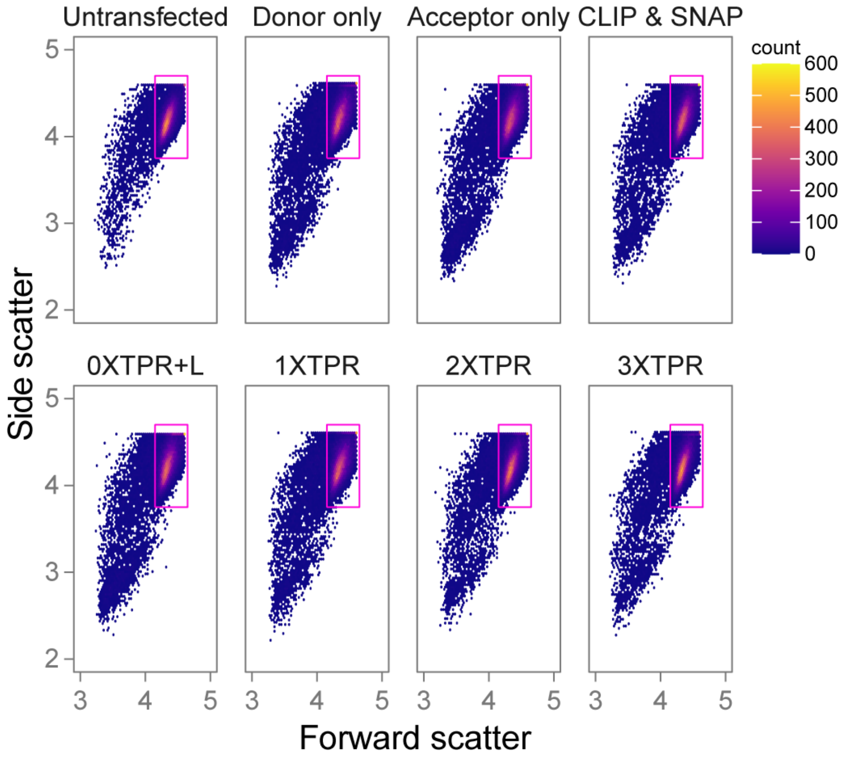


**Figure S5: Gating of single live cells from flow cytometry based-FRET.**
Hexbin plot of forward and side scatter for untransfected cells (n = 27,026), 0XTPR+L donor only (n = 33,758), 0XTPR+L acceptor only (n = 33,588), CLIP & SNAP cotransfection (n = 33,185), 0XTPR+L (n = 34,177), 1XTPR (n = 33,431), 2XTPR (n = 30,764) and 3XTPR (n = 31,008). Gate to select single live cells is shown in magenta..

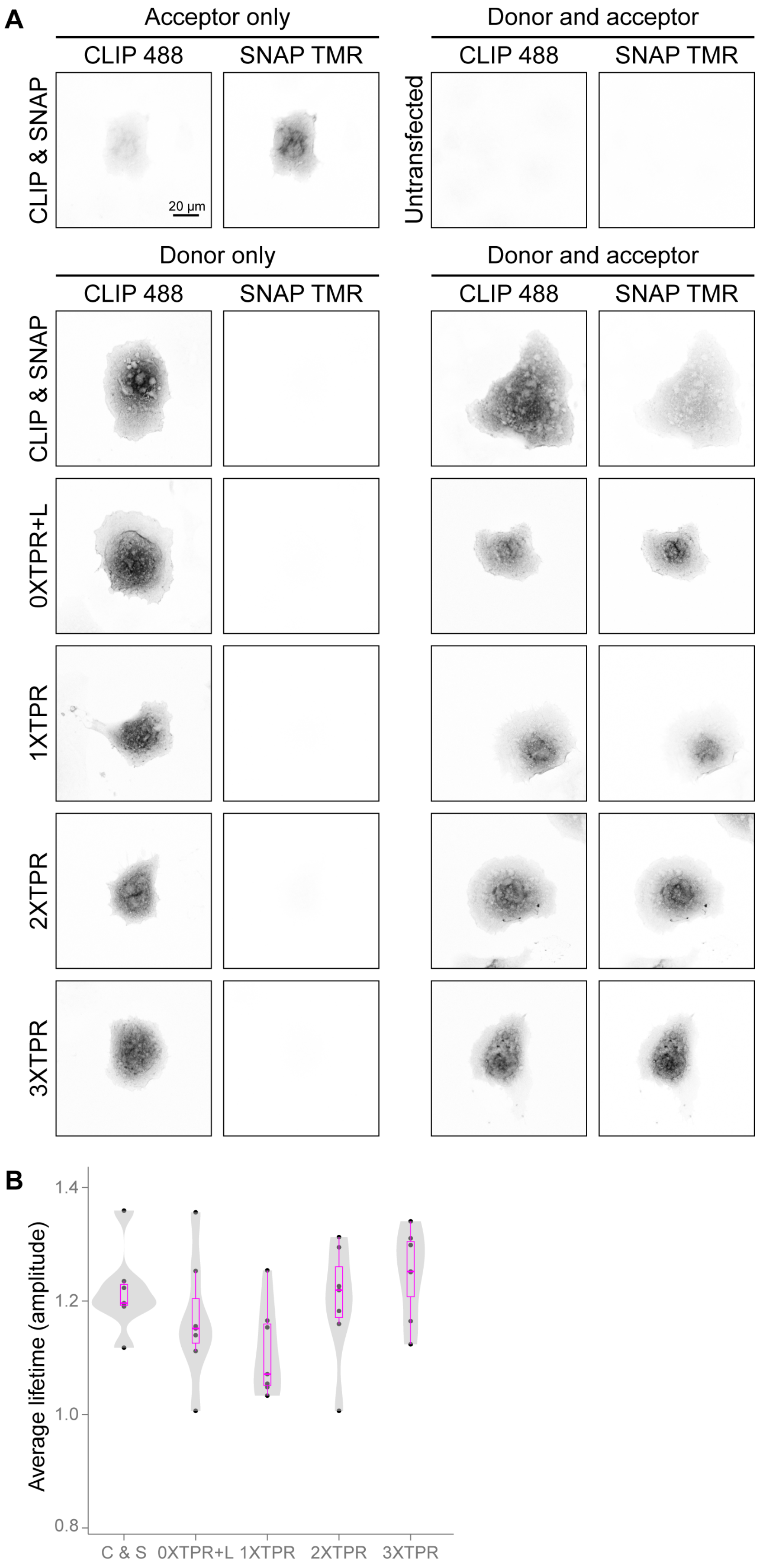


**Figure S6: Labelling of COS-7 cells for FLIM-FRET.**

(A) Representative confocal images of COS-7 cells cotransfected with CLIP and SNAP, transfected with one of the four FRET standards (0XTPR+L, 1XTPR, 2XTPR, 3XTPR) or untransfected. Cells were labelled with; SNAP-Cell TMR-Star (acceptor only, top left), CLIP-Surface 488 (donor only, bottom left) or both fluorophores (donor and acceptor, right). Images show the labelling and lack of bleed through between channels for the donor.

(B) Average lifetime (amplitude) of the four FRET standards; 0XTPR+L (n = 7), 1XTPR (n = 7), 2XTPR (n = 7), 3XTPR (n = 7) or a cotransfection of CLIP and SNAP (n = 7) for samples labelled with only CLIP-Surface 488 (donor only) from (6C). Results from the FRET standards were compared to the CLIP SNAP cotransfection by a one-way ANOVA and all were not significantly different.
